# Carotenoid retention during post-harvest storage of *Capsicum annuum*: the role of the fruit surface structure

**DOI:** 10.1101/2023.01.27.525890

**Authors:** Alexandra C. Holden, Hagai Cohen, V. Rickett Daniel, Asaph Aharoni, Paul D. Fraser

**Author notes:** Department of vegetable and field crops, Institute of Plant Sciences, ARO, Volcani Center, 68 HaMaccabim Rd, Rishon, LeZion, 7505101. Israel.

## Abstract

Degradation of carotenoids in food crops during post-harvest storage results in major economic and nutritional losses. In this study, a pepper (*Capsicum annuum*) panel for post-harvest carotenoid retention was studied to elucidate underlying mechanisms associated with this commercial trait of interest. Quantitative determination of carotenoid pigments and concurrent cellular analysis indicated that those pepper fruit with thicker lipid exocarp layers and smooth surfaces, following post-harvest drying and storage, possessed increased carotenoid retention. Total cutin monomer content increased in high carotenoid retention fruits and sub-epidermal cutin deposits were responsible for the difference in exocarp thickness. Cutin biosynthesis and cuticle precursor transport genes were differentially expressed between high and low carotenoid retention genotypes, and this supports the finding that fruit cuticle biosynthesis is associated with carotenoid retention. Carotenoids were located within cells embedded within the sub-epidermal cutin layer, and these carotenoids were protected from degradation due to the lack of permeability of the fruit surface to reactive oxygen species, and their precursors. The identification of a novel role for the pepper fruit surface in protecting against carotenoid degradation serves as an important discovery for the function of the fruit cuticle and provides an exploitable resource to enhance fruit quality.

**Highlight statement:** Carotenoid pigments in Chilli pepper confer post-harvest colour and nutritional quality. Analysis of diverse commercial genotypes indicates the involvement of the fruit surface in carotenoid retention

## Introduction

The red, yellow, and orange colours observed in pepper (*Capsicum annuum*) fruits is conferred by the presence of carotenoids. Brightly coloured pepper fruits, with uniform colour distribution, are of high quality to consumers. Along with the colour profile of freshly harvested fruits, the ability of pepper fruits to retain colour for long periods post-harvest is an equally important pepper fruit quality trait. The pepper crop is in demand from consumers year-round, however, due to environmental conditions in pepper growing regions, the crop cannot be grown and harvested throughout the year. Therefore, storage of dry peppers following harvest is essential. Pepper fruits must retain quality during post-harvest storage, including colour. Post-harvest storage losses are not only observed in fruit crops, but in other important crops, including rice, wheat, and maize (Kumar and Kalita, 2017). Carotenoid losses during post-harvest storage are reported in a variety of crop species (Burt et al., 2010, Bechoff et al., 2011), and this affects colour and nutritional quality. Furthermore, whilst increased provitamin A capacity has been engineered in rice, resulting in a carotenoid enriched variety known as ‘Golden rice’ (Beyer et al., 2002), maintaining such increased nutritional properties has proved challenging, as carotenoid degradation occurs upon post-harvest storage (Gayen et al., 2015).

Carotenoids are lipophilic tetraterpenes, with a conjugated double bond structure, which results in the formation of a chromophore. The length of chromophore determines the absorption spectrum of the molecule, and hence, the perceived colour (Fraser and Bramley, 2004). Capsanthin and capsorubin are the two major carotenoids found in red pepper fruits, which are almost unique to the ripe fruit of peppers. Capsanthin and capsorubin are synthesised from antheraxanthin and violaxanthin, respectively, by the action of CAPSANTHIN/CAPSORUBIN SYNTHASE (CCS) (Bouvier et al., 1994). Pepper carotenoids are commonly found to be esterified (Biacs et al., 1989, Minguez-Mosquera and Hornero-Mendez, 1994b), making them more stable than free carotenoids (Biacs et al., 1989). Carotenoids not only confer colour properties to pepper fruits, but further have high antioxidant capacity and are therefore beneficial to human health. This antioxidant capacity is a result of their ability to scavenge reactive oxygen species, due to their conjugated double bond structure. Upon interaction of carotenoids with singlet oxygen (^1^O_2_), physical quenching occurs, in which energy is transferred between the two molecules. The excess energy of ^1^O_2_ is transferred to the carotenoid and yields ground state oxygen and a triplet excited carotene (Stahl and Sies, 2003). Oxidation of carotenoids results in the formation of cleavage products, collectively known as apocarotenoids. Whilst apocarotenoids have critical functions within the cell, including as photooxidation protection, signalling molecules, and phytohormone precursors (Hou et al., 2016), excessive carotenoid degradation results in colour loss.

Waxy cuticles are an essential part of a plant’s physiology and play a fundamental role in the plant’s interaction with the environment, including mitigating the negative effects of excessive water loss (Domínguez et al., 2011). The cuticle has been further suggested to have a role in the post-harvest quality of fruit (Lara et al., 2014, Lara et al., 2019). Plant cuticles are comprised of two components: cutin monomers and waxy, or lipophilic components, which are embedded within the cutin matrix, as reviewed in Yeats and Rose, 2013. The cuticle provides a barrier to control the entrance and exit of gases in fruits, which lack stomata. Intact pepper fruit cuticle has been shown to be permeable to a small amount of carbon dioxide and oxygen, though this permeability increases significantly upon wounding of the cuticle (Banks and Nicholson, 2000). Whilst gaseous exchange is essential, negative effects may be observed as a result. The role of antioxidants is evident in protecting against the harmful effects of reactive oxygen species, which may be formed during the process of gaseous exchange.

Pepper fruit cuticle has been reported to extend through the apoplast of multiple cell layers (Martin and Rose, 2014). There is some dispute over whether this ‘cuticular’ layer, which penetrates several cell layers deep, can be termed the ‘cuticle’. For this reason, the ‘cuticular layer’ spanning several cell layers, has been termed the ‘wax exocarp’, which is defined as the outermost lipophilic layer of the pericarp of an angiosperm fruit, external to the mesocarp. Pepper fruit cuticular components, and associated cuticle morphology QTLs have been previously identified (Popovsky-Sarid et al., 2017).

In the present study, the influence of the pepper fruit cuticle and surface structure on the carotenoid retention trait is described, with the cuticle playing a role in protecting carotenoids from degradation, particularly during post-harvest storage. In addition, the implications for this novel finding for engineering carotenoid enriched crops and for breeding desirable traits into pepper are discussed.

## Results

### Carotenoid retention classification of the commercial panel

A commercial panel, consisting of 13 independent pepper genotypes displaying variation in their carotenoid content and carotenoid retention phenotype, was grown, oven dried, and stored in a manner to replicate commercial conditions. HPLC analysis was used to quantify carotenoid content of pepper fruits before oven drying, and subsequently throughout the post-harvest storage period. Carotenoid retention was then calculated. Over this period, the 13 genotypes displayed varying levels of carotenoids: genotype R3 contained only 2.5 mg/g dry weight of carotenoids at the fresh fruit stage, whereas genotype R7 contained more than 18 mg/g dry weight of carotenoids at the same fruit stage (*Table S1*). Furthermore, the change in carotenoid content during the twelve week post-harvest storage period differed between the 13 pepper varieties analysed. Carotenoid retention was calculated as the change in carotenoid content between the fresh fruit time point, and following 12 weeks of post-harvest storage. Some genotypes displayed very little change in total carotenoid content during post-harvest 4 °C storage, such as lines R1 and R12, whereas other lines showed significant increases (R5) or decreases (R7) in total carotenoid content.

Pepper genotypes were characterised as low, medium, or high carotenoid retention dependent on the change in carotenoid content during post-harvest storage. Arbitrary values were used to determine carotenoid retention phenotypes: lines which decreased in carotenoid content by more than 10% were deemed to be low retention (R3, R4), lines which showed a change in carotenoid content between -10% and 10% were deemed to be medium carotenoid retention (R1, R2), and lines which increased in carotenoid content by more than 10% were characterised as high retention (R5, R6).

Interestingly, carotenoid retention phenotypes were assigned previously using a visual method, however, carotenoid quantification was deemed to be a more accurate method for characterising carotenoid retention.

### Fruit surface texture is associated with carotenoid retention following post-harvest storage

Post-harvest drying of pepper fruits resulted in vastly different fruit surface textures when comparing the 13 pepper genotypes. Following oven drying, some pepper fruits, such as genotypes R1, R6, and R8, retained the smooth, waxy surface texture that all genotypes had at the fresh time point, whilst other genotypes, such as R3, R4, and R7, displayed significant surface ‘wrinkling’. Therefore, all pepper genotypes were characterised as having either a smooth cuticle surface or a cracked cuticle surface (*Fig. S1*). Genotypes characterised as low carotenoid retention tended to display a wrinkled surface texture following drying, whilst medium and high carotenoid retention fruits tended to display a smooth surface texture following drying.

Scanning electron microscopy (SEM) further supported these differences between high and low carotenoid retention pepper genotypes in their fruit surface texture. Low carotenoid retention genotypes, R3 and R4, both had a ‘wrinkled’ surface texture, and showed evidence of surface cracks (Fig. 1B), whilst medium and high carotenoid retention genotypes, R6 and R8, both had smooth surface textures with no cracks present on the surface.

**Fig. 1.**
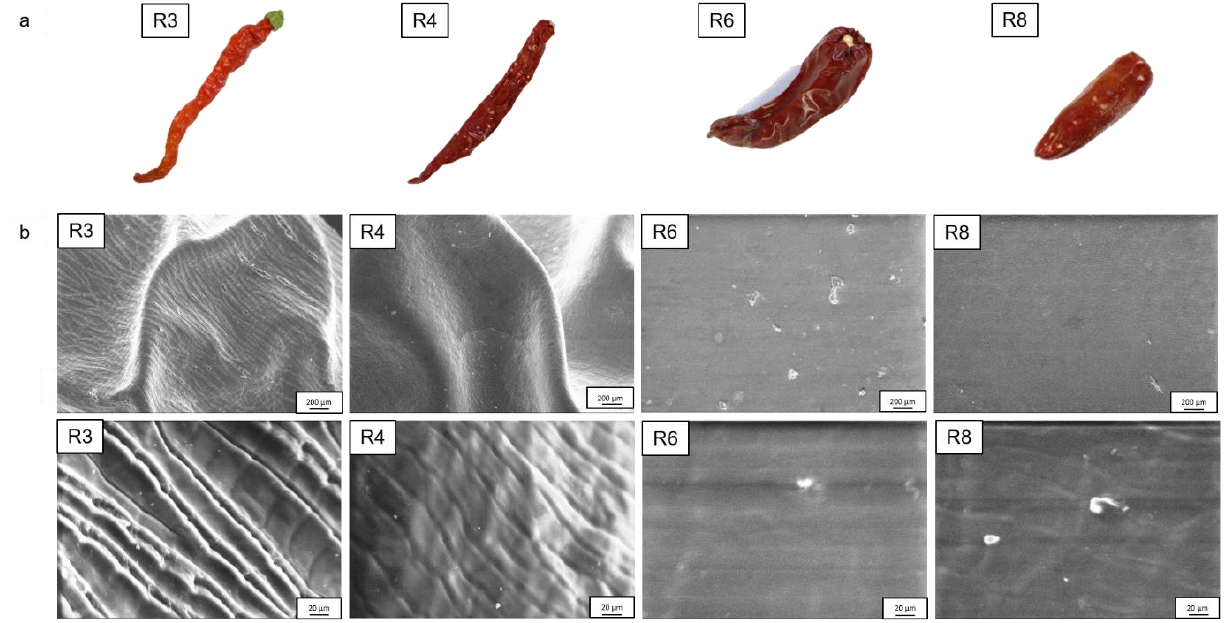
Fruit surface texture is associated with the carotenoid retention trait following post-harvest storage. Panel a: Pepper fruits following oven drying. R3 and R4 display ‘wrinkled’ surface texture after drying; R6 and R8 display ‘smooth’ surface texture after drying. Panel b: Scanning electron microscopy images of dried pepper fruit surface. R3 and R4 display surface cracks; R6 and R8 display smooth surfaces. 300x and 3000x magnifications used.

### Fruit exocarp thickness is associated with the carotenoid retention phenotype

Due to the differences observed between high and low carotenoid retention genotypes in fruit surface structure following post-harvest drying, further analysis of the fruit surface structure was carried out.

The outer surface of the pepper fruit, defined as the fruit exocarp, of fruits within the pepper diversity panel were observed using light microscopy, in order to determine whether variation in the fruit exocarp structure is associated with the carotenoid retention phenotype. Nile Red was used to stain the wax exocarp and Fast Green was used to stain pericarp, in order to clearly differentiate between these two tissue types. This method demonstrated that there was a clear difference in exocarp thickness within the fruits of the 13 pepper genotypes studied (Fig. 2A; Fig. S2). Staining showed that epidermal cells were embedded within the wax exocarp layer, and in some cases, the wax exocarp layer penetrated several cell layers deep into the fruit. Whilst some genotypes had a very thin exocarp layer, for example R3 and R7, which showed just one layer of cells embedded within the wax exocarp layer, other genotypes, such as R5 and R6, displayed up to four or five layers of cells embedded within the wax exocarp. This difference in exocarp thickness was further supported following cryo-SEM, which demonstrated that low carotenoid retention lines R3 and R4 both had just one layer of cells embedded within the outer layer, whilst lines R6 and R8 both displayed several cell layers embedded within the outer layer.

**Fig. 2.**
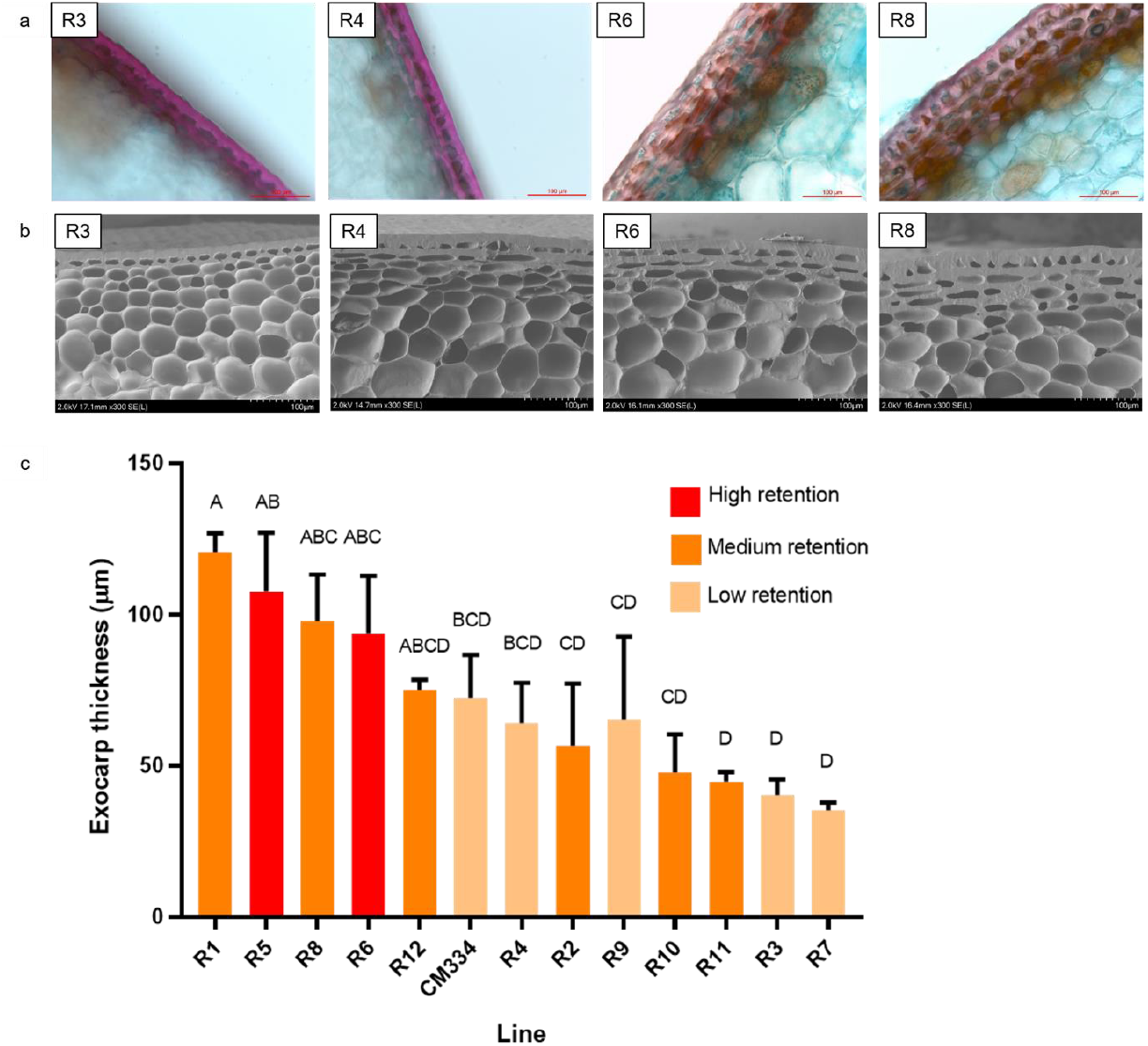
Exocarp thickness is associated with the carotenoid retention phenotype. Panel a: Light microscopy images of pepper fruit exocarp. Panel b: Cryo scanning electron microscopy images of pepper fruit exocarp. Panel c: Exocarp thickness of pepper diversity panel measured using light microscopy images. Mean ± SEM bars reported.

Clear differences were observed in the thickness of the wax exocarp layer between different genotypes within the pepper diversity panel. Whilst lines R1, R5, R6, and R8 all had an exocarp thickness ranging from 90-120 µm, lines R3 and R7 both had a thinner exocarp, with a thickness between 30-40 µm. The general trend appeared in which genotypes characterised as high or medium carotenoid retention also tended to have a thicker exocarp.

### Biochemical profiling of pepper fruit cuticle

The carotenoid retention phenotype is only observed upon ripening of the pepper fruit, cuticle composition was analysed in ripe fruit using an established GC-MS-based method, as this was deemed to be the most biologically relevant developmental stage for this trait. As a comparator, the cuticle of mature green fruit was also profiled (*Table S2*).

Significantly lower levels of 10,16-dihydroxyhexadecanoic acid were observed in both low carotenoid retention genotypes, R3 and R4, when compared to the medium/high carotenoid retention genotype, R8, however not compared to R6 (Fig. 3E). This compound accounts for approximately 90% of the total content of the cutin polymer, suggesting that low carotenoid retention is associated with lower total cutin content. In ripe fruit, R3 had significantly lower levels of ferulic acid compared to genotypes R4 and R6 (Fig. 3A), and octadeca-9,12-dienoate compared to genotypes R4, R6, and R8 (Fig. 3C). The low carotenoid retention genotype, R3, displayed significantly lower total cutin monomer content when compared to the medium/high carotenoid retention genotype, R8, but no significant difference was observed between other genotypes (Fig. 3G). Overall, the differences in cutin monomer content reflect the differences observed in pepper exocarp thickness. The low carotenoid retention genotype, R3, displayed a significantly thinner exocarp compared to the medium/high carotenoid retention genotype, R8, and this significant difference between these two genotypes is further supported by the differences in their cutin compositions. The high carotenoid retention genotype, R6, also showed a significantly thicker exocarp compared to R3, however no difference in total cutin monomer was reported between these two varieties. Consequently, only a general trend is evident, in which increased cutin monomer content was observed in genotypes displaying high carotenoid retention phenotypes.

**Fig. 3.**
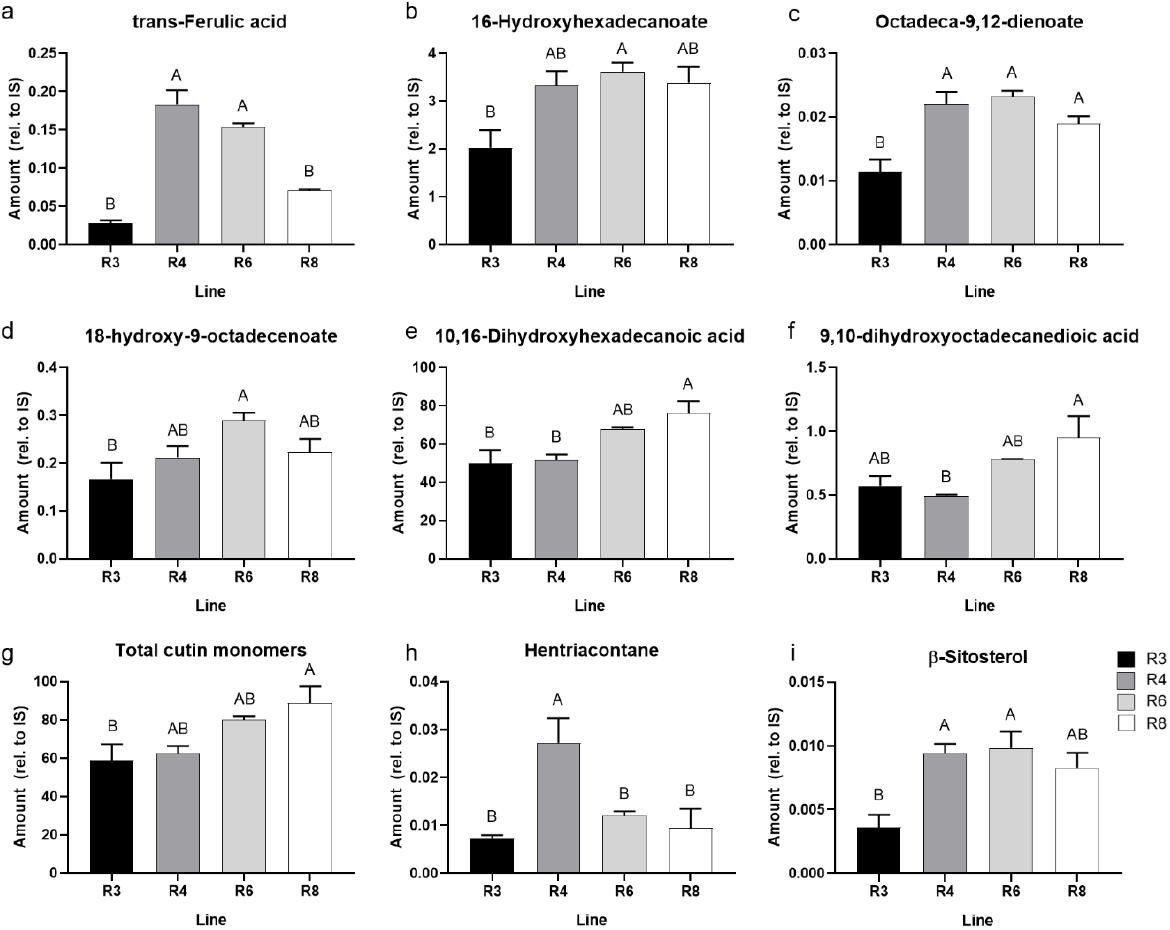
Cuticle components of ripe fruit associated with the carotenoid retention phenotype. Cutin monomer components and cuticular waxes were measured in ripe fruits displaying variation in carotenoid retention phenotype: R3 = low carotenoid retention, R4 = low carotenoid retention, R6 = high carotenoid retention, R8 = medium/high carotenoid retention. Panel a-g: cutin monomer components, Panel h, i: cuticular lipophilic components. Only compounds in which significant differences were reported between pepper genotypes are reported here. Mean ± SEM bars reported. C_32_ alkane (100 µg) was utilised as an internal standard in cutin monomer extraction; Deuterated traiacontane (C_30_) (10 µg) was utilised as an internal standard in cuticle lipophilic component extraction. Amounts were calculated as relative to respective internal standards.

Cuticular lipophilic components, which are embedded within the cutin matrix, were analysed in the four pepper fruit genotypes (R3, R4, R6, R8). Whilst β-sitosterol levels were decreased in the low carotenoid retention genotype, R3 (Fig. 3I), no other significant differences, in which cuticular lipophilic composition and cuticle thickness were associated, were detected (Table S2). The low carotenoid retention genotype, R4, displayed significantly higher levels of hentriacontane compared to the other genotypes analysed (Fig. 3H), however, this did not follow the emerging trend in which exocarp thickness, and hence cuticle composition, associates with the carotenoid retention phenotype. Consequently, cuticle lipophilic components did not appear to be associated with the carotenoid retention phenotype, nor with exocarp thickness.

### Cuticle biosynthesis gene expression displays an association with changes observed in cuticle structure between high and low carotenoid retention pepper genotypes

RNAseq was used to determine gene expression patterns in pepper fruit epidermal cells during cuticle development. RNA was isolated from pepper fruit epidermal cells of genotypes R3 and R8, in order to provide spatiotemporal resolution to gene expression analyses. Genes involved in cuticle biosynthesis were identified in pepper, based on homology with orthologs in other species, which have been reported to be involved in cuticle biosynthesis. Differentially expressed genes between the high and low carotenoid retention lines are reported in Table 2. The pepper ortholog of the ABC transporter PERMEABLE CUTICLE1/ABCG32 (PEC1), which functions in the transfer of cutin components over the plasma membrane in *Arabidopsis thaliana* (Bessire et al., 2011), was identified as significantly up-regulated in the high carotenoid retention genotype compared to the low carotenoid retention genotype. Lipid transport proteins (LTPs) are considered to be involved in the transport of cutin monomers to the site of cuticle synthesis (Yeats and Rose, 2008). Several LTPs were identified as differentially expressed between the high and low carotenoid retention genotypes, however some were up-regulated in the high retention genotype, whilst others were down-regulated. *Bodyguard* (*BDG*) has been identified as playing a role in cutin synthesis (Kurdyukov et al., 2006), and an ortholog of this gene was found to be up-regulated in the high carotenoid retention genotype compared to the low retention genotype. Interestingly, a MYB16 transcription factor was found to be down-regulated in the high carotenoid retention genotype early in fruit development. The ortholog of this gene has been shown to be involved in the regulation of cuticle development, along with *WIN1/SHN1* in *Arabidopsis thaliana* (Oshima et al., 2013), and so it may have been expected that this gene would be up-regulated in the high carotenoid retention genotype, as these fruits had a thicker fruit cuticle compared to the low carotenoid retention genotype. However, *WIN1/SHN1* orthologs were not found to be differentially expressed in pepper. Two genes were annotated as *CER1* in the pepper Sol Genomics Network, and interestingly they displayed differences in expression between the high and low retention genotypes. CA09G18740 was shown to be up-regulated in the high retention line at anthesis + 45 days, whilst CA12G22670, also annotated as *CER1*, was shown to be down-regulated in the high retention genotype in ripe fruits. *CER1* is involved in reduction and decarbonylation of the very long chain fatty acids to cuticular alkanes (Bernard et al., 2012). A cytochrome P450 enzyme MAH1, a midchain alkane hydroxylase, is involved in the oxidation of alkanes to secondary alcohols and ketones in *Arabidopsis* (Greer et al., 2007). *MAH1* genes were found to be up-regulated in the high carotenoid retention genotypes compared to the low carotenoid retention genotype throughout fruit development (Table 2).

**Table 1:** Change in carotenoid content following post-harvest storage of diverse pepper genotypes. Carotenoid content measured before and after post-harvest storage, and change in carotenoid content calculated. Carotenoid retention classification allocated based on change in carotenoid content.

| Pepper genotype | Change in carotenoid content (%) | Carotenoid retention |
| --- | --- | --- |
| R1 | -0.6 | Medium |
| R2 | 2.7 | Medium |
| R3 | -18.3 | Low |
| R4 | -20.9 | Low |
| R5 | 48.3 | High |
| R6 | 24.2 | High |
| R7 | -25.7 | Low |
| R8 | -0.6 | Medium |
| R9 | -17.3 | Low |
| R10 | 8.5 | Medium |
| R11 | 5.7 | Medium |
| R12 | -1.0 | Medium |
| CM334 | -18.0 | Low |

**Table 2:**
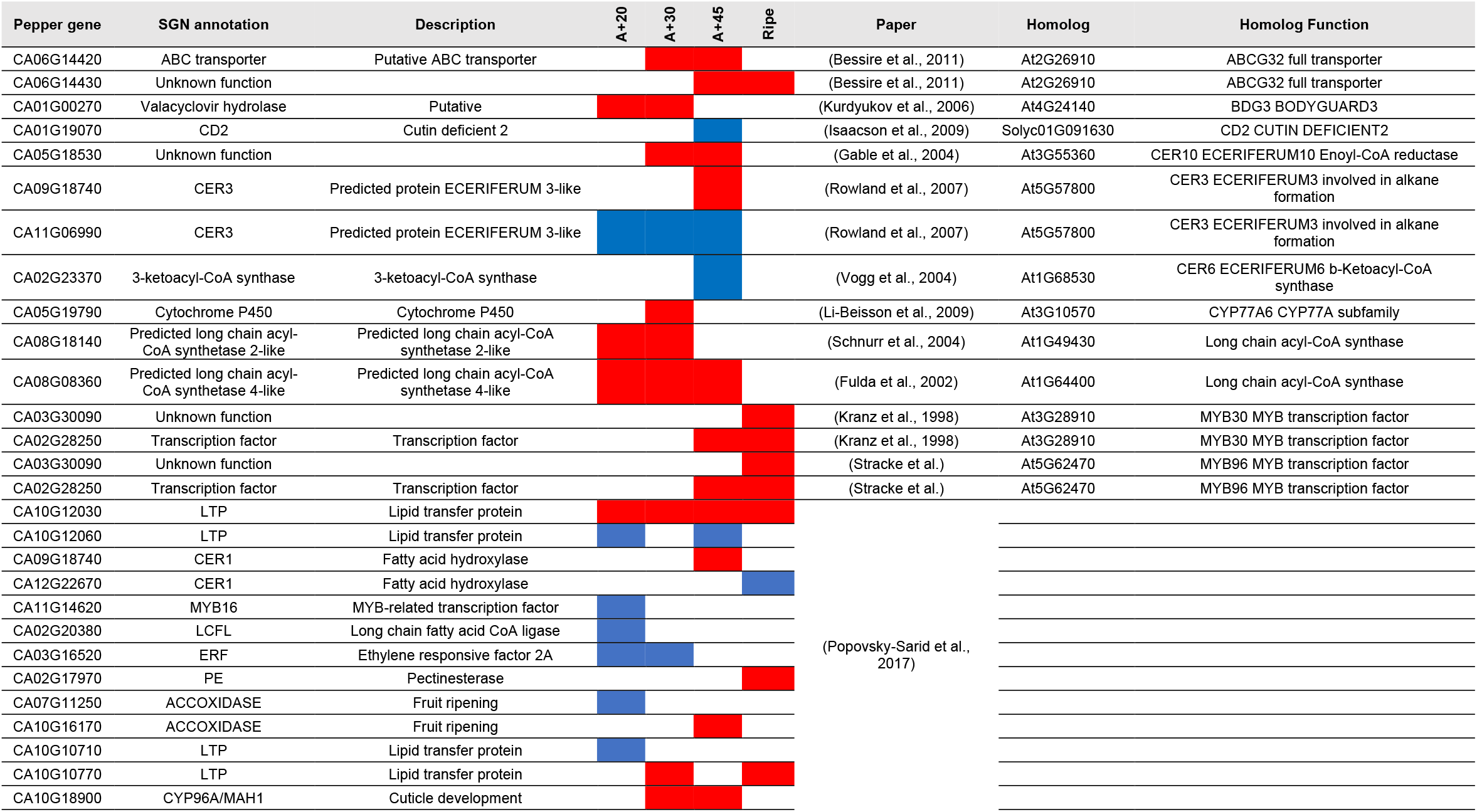
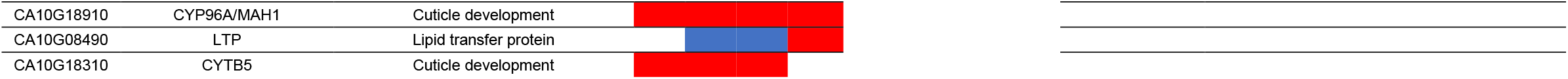
Differentially expressed genes associated with fruit cuticle biosynthesis. Red = down regulated R3, Blue = up regulated R3. Transcript number was assessed at four fruit development stages: anthesis + 20 days, anthesis + 30 days, anthesis + 45 days, and ripe. Genes were identified based on literature searches, and using BLAST to identify gene orthologs. Sol Genomic Network gene descriptions are noted, if provided. Genes were considered to be significantly differentially expressed if p < 0.05, FDR < 0.05.

| Pepper gene | SGN annotation | Description | A+20 | A+30 | A+45 | Ripe | Paper | Homolog | Homolog Function |
| --- | --- | --- | --- | --- | --- | --- | --- | --- | --- |
| CA06G14420 | ABC transporter | Putative ABC transporter |  |  |  |  | (Bessire et al., 2011) | At2G26910 | ABCG32 full transporter |
| CA06G14430 | Unknown function |  |  |  |  |  | (Bessire et al., 2011) | At2G26910 | ABCG32 full transporter |
| CA01G00270 | Valacyclovir hydrolase | Putative |  |  |  |  | (Kurdyukov et al., 2006) | At4G24140 | BDG3 BODYGUARD3 |
| CA01G19070 | CD2 | Cutin deficient 2 |  |  |  |  | (Isaacson et al., 2009) | Solyc01G091630 | CD2 CUTIN DEFICIENT2 |
| CA05G18530 | Unknown function |  |  |  |  |  | (Gable et al., 2004) | At3G55360 | CER10 ECERIFERUM10 Enoyl-CoA reductase |
| CA09G18740 | CER3 | Predicted protein ECERIFERUM 3-like |  |  |  |  | (Rowland et al., 2007) | At5G57800 | CER3 ECERIFERUM3 involved in alkane formation |
| CA11G06990 | CER3 | Predicted protein ECERIFERUM 3-like |  |  |  |  | (Rowland et al., 2007) | At5G57800 | CER3 ECERIFERUM3 involved in alkane formation |
| CA02G23370 | 3-ketoacyl-CoA synthase | 3-ketoacyl-CoA synthase |  |  |  |  | (Vogg et al., 2004) | At1G68530 | CER6 ECERIFERUM6 b-Ketoacyl-CoA synthase |
| CA05G19790 | Cytochrome P450 | Cytochrome P450 |  |  |  |  | (Li-Beisson et al., 2009) | At3G10570 | CYP77A6 CYP77A subfamily |
| CA08G18140 | Predicted long chain acyl-CoA synthetase 2-like | Predicted long chain acyl-CoA synthetase 2-like |  |  |  |  | (Schnurr et al., 2004) | At1G49430 | Long chain acyl-CoA synthase |
| CA08G08360 | Predicted long chain acyl-CoA synthetase 4-like | Predicted long chain acyl-CoA synthetase 4-like |  |  |  |  | (Fulda et al., 2002) | At1G64400 | Long chain acyl-CoA synthase |
| CA03G30090 | Unknown function |  |  |  |  |  | (Kranz et al., 1998) | At3G28910 | MYB30 MYB transcription factor |
| CA02G28250 | Transcription factor | Transcription factor |  |  |  |  | (Kranz et al., 1998) | At3G28910 | MYB30 MYB transcription factor |
| CA03G30090 | Unknown function |  |  |  |  |  | (Stracke et al.) | At5G62470 | MYB96 MYB transcription factor |
| CA02G28250 | Transcription factor | Transcription factor |  |  |  |  | (Stracke et al.) | At5G62470 | MYB96 MYB transcription factor |
| CA10G12030 | LTP | Lipid transfer protein |  |  |  |  | (Popovsky-Sarid et al., 2017) |  |  |
| CA10G12060 | LTP | Lipid transfer protein |  |  |  |  |  |  |  |
| CA09G18740 | CER1 | Fatty acid hydroxylase |  |  |  |  |  |  |  |
| CA12G22670 | CER1 | Fatty acid hydroxylase |  |  |  |  |  |  |  |
| CA11G14620 | MYB16 | MYB-related transcription factor |  |  |  |  |  |  |  |
| CA02G20380 | LCFL | Long chain fatty acid CoA ligase |  |  |  |  |  |  |  |
| CA03G16520 | ERF | Ethylene responsive factor 2A |  |  |  |  |  |  |  |
| CA02G17970 | PE | Pectinesterase |  |  |  |  |  |  |  |
| CA07G11250 | ACCOXIDASE | Fruit ripening |  |  |  |  |  |  |  |
| CA10G16170 | ACCOXIDASE | Fruit ripening |  |  |  |  |  |  |  |
| CA10G10710 | LTP | Lipid transfer protein |  |  |  |  |  |  |  |
| CA10G10770 | LTP | Lipid transfer protein |  |  |  |  |  |  |  |
| CA10G18900 | CYP96A/MAH1 | Cuticle development |  |  |  |  |  |  |  |

|  |  |  |
| --- | --- | --- |
| CA10G18910 | CYP96A/MAH1 | Cuticle development |
| CA10G08490 | LTP | Lipid transfer protein |
| CA10G18310 | CYTB5 | Cuticle development |

As genes involved in cuticle biosynthesis have been shown to be differentially expressed between the high and low carotenoid retention genotypes, this provides further evidence that the carotenoid retention phenotype is controlled, in at least some part, by the structure of the fruit surface.

### Carotenoids associated with the fruit exocarp may influence the carotenoid retention trait

Upon enzymatically isolating exocarp discs from pepper fruit, it was unexpectedly noted that red pigment was retained within the wax exocarp tissue, despite the removal of all other pericarp tissue (Fig. 4A). Consequently, it was hypothesised that carotenoids may be located within the epidermal cells found in the isolated exocarp layer. As the waxy exocarp layer penetrated several cell layers deep within the fruit, particularly in genotypes R6 and R8, these carotenoids may still remain within the exocarp due to the protective nature of the exocarp layer. Comparatively, upon isolation of tomato fruit exocarp discs, the red pigment observed in pepper exocarp discs was not present in tomato discs, and further, upon washing solvent extraction, pigment was not removed from tomato exocarp discs in the same manner as to which it was isolated in pepper exocarp discs (Fig. S3).

**Fig. 4.**
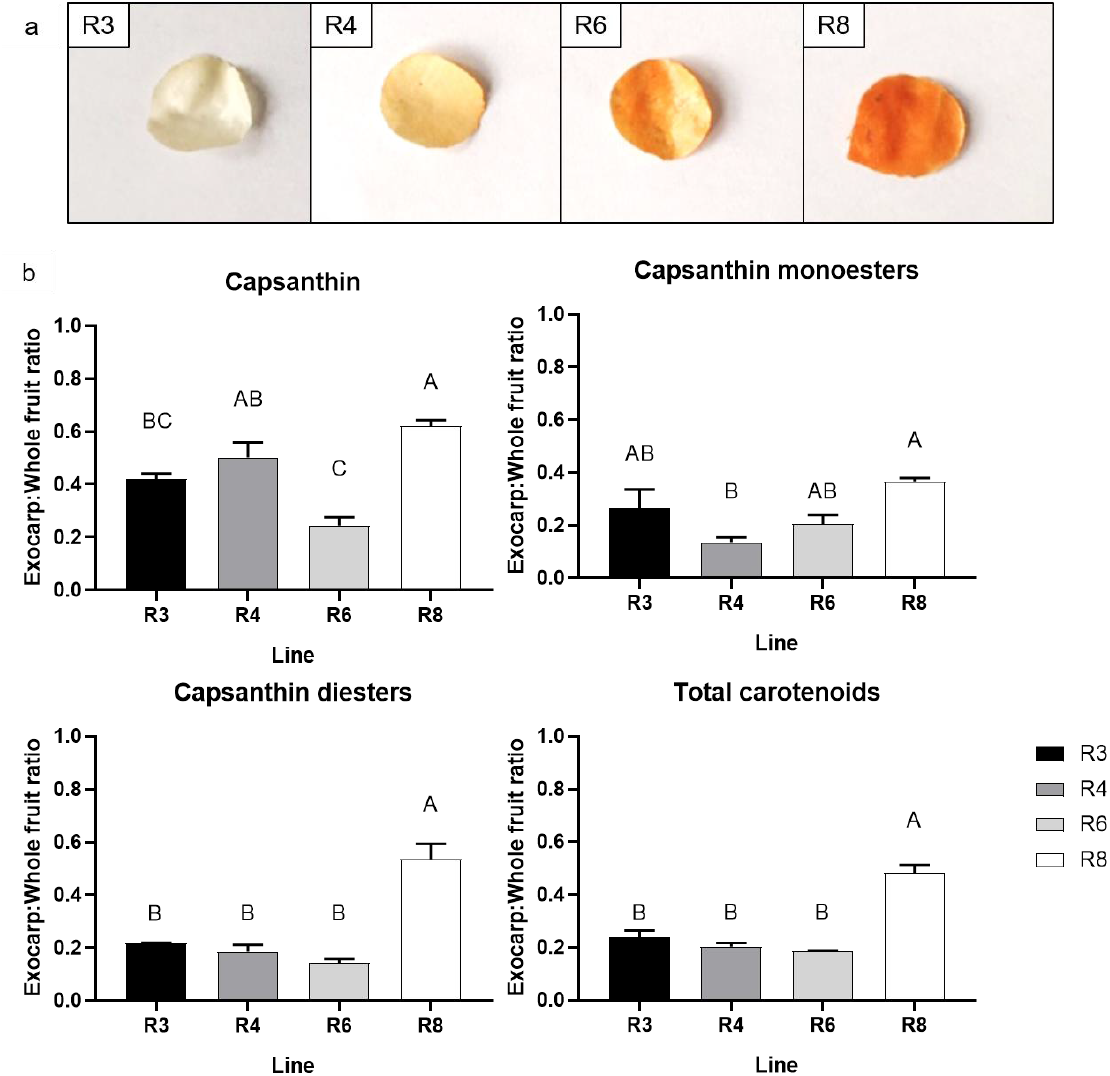
Carotenoids are associated with pepper fruit exocarp. Panel a: Pepper fruit exocarp discs (1 cm diameter). Panel b: Carotenoid content of pepper exocarp discs, expressed as exocarp to whole fruit ratio. Mean ± SEM bars reported.

Carotenoids were isolated from exocarp discs through a process of washing in chloroform and methanol over a period of three days. HPLC analysis was used to quantify carotenoids extracted from exocarp discs. These values were then compared to carotenoid amounts extracted from whole fruit discs, and results were expressed as a ratio (Figure 4B). This approach allowed the quantity of exocarp-bound carotenoids to be compared in a relative manner to whole fruit carotenoid content, across all genotypes analysed, regardless of variation in total fruit carotenoid content between genotypes. Interestingly, significant differences were only observed in the ratio of capsanthin and capsanthin esters between the genotypes analysed. This suggests that these carotenoids may be responsible for causing the observed red colour of exocarp discs in these genotypes. A significantly greater exocarp to whole fruit ratio for capsanthin diesters was observed in the high carotenoid retention genotype, R8, relative to the three other genotypes analysed. This trend was further reflected in total carotenoid content when comparing R8 to the other genotypes analysed.

### Initiation of carotenoid degradation reveals crucial protective role of exocarp

The fruit surface structure has been demonstrated to be associated with the carotenoid retention phenotype, it was hypothesised that the fruit surface may be crucial in protecting against carotenoid degradation. Cracks were not observed on the surface of high carotenoid retention fruits; therefore, it could be postulated that the fruit surface may provide a protective barrier against oxidative degradation of carotenoids.

Pepper fruits were treated with hydrogen peroxide, as an oxidative agent, in order to determine the role of the fruit surface in protecting against carotenoid degradation. Fruits were harvested when ripe, before control fruits were treated with varying concentrations of hydrogen peroxide immediately following harvest, and were oven dried, and stored for four weeks.

Following post-harvest storage, a decrease in total carotenoid content was observed in the low carotenoid retention genotype, R3, peppers treated with 2 mM H_2_O_2_ before drying when compared to peppers treated with 0 mM H_2_O_2_ (Fig. 5A), at which point, the cuticle remained smooth and exhibited no cracks. No decrease was observed in peppers treated with 0.2 mM H_2_O_2_ before drying. In contrast, R3 peppers treated with both 0.2 mM and 2 mM H_2_O_2_ after fruit drying displayed decreases in total carotenoid content (Fig. 5B) compared to peppers treated with 0 mM H_2_O_2_ at this time point. Fruits treated with H_2_O_2_ following drying were more susceptible to carotenoid degradation, when a lower concentration (0.2 mM) H_2_O_2_ was used. After drying the fruit surface displayed cracking, this structural alteration may have allowed the entrance of oxidative species into the fruit and promoted carotenoid degradation.

**Fig. 5.**
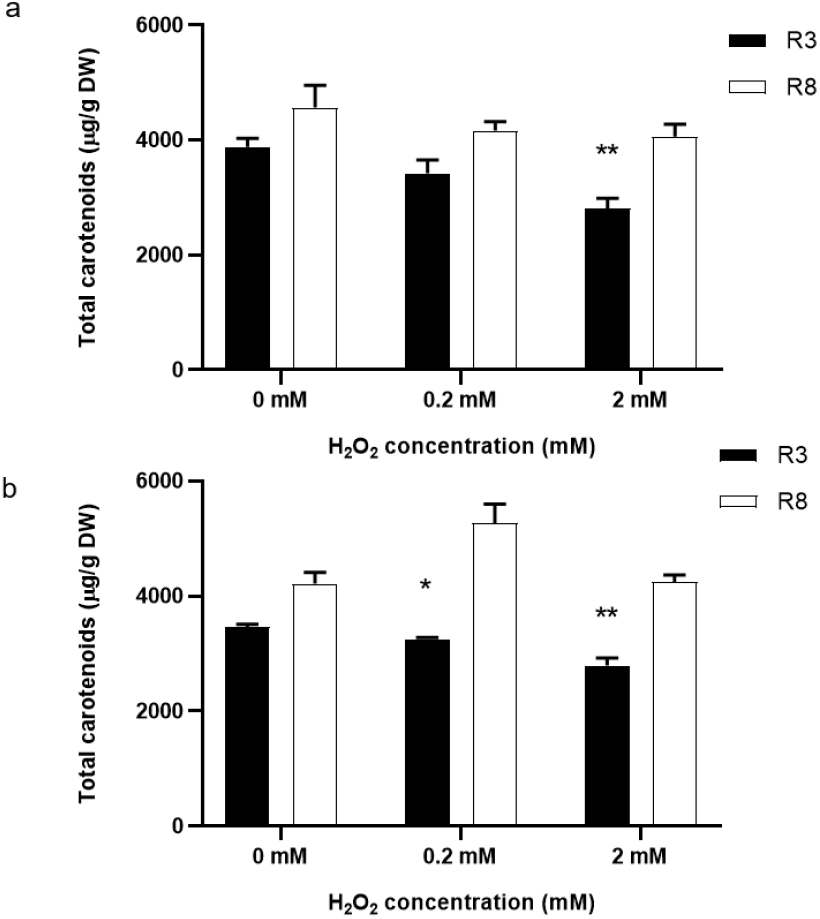
Carotenoid content of pepper fruits after post-harvest storage, following treatment with hydrogen peroxide. Panel a: Total carotenoid content (µg/g DW) after post-harvest storage, following hydrogen peroxide treatment before fruit drying. Panel b: Total carotenoid content (µg/g DW) after post-harvest storage, following hydrogen peroxide treatment after fruit drying. Student’s t test used to determine significant differences between the control condition (0 mM H_2_O_2_) and test concentrations. Mean ± SEM bars reported.

The medium/high carotenoid retention genotype, R8, displayed no difference in total carotenoid content, regardless of H_2_O_2_ concentration used (Fig. 5A, B). This was observed in fruits treated both before and after fruit drying. As the fruit surface remained smooth and intact following drying, a protective barrier was in place to limit the entry of oxidative species into the fruit. Consequently, no difference in carotenoid content was observed, even when relatively high concentrations of H_2_O_2_ (2 mM) were used.

## Discussion

### Carotenoid retention as a key quality trait of pepper fruits

Pepper fruit colour is important as a quality trait immediately following harvest from the plant. However, fruits may be dried and stored for several months in industry, and therefore, pepper colour following post-harvest storage is an equally important quality trait. The major carotenoid conferring red colour in chilli peppers is capsanthin, whilst other xanthophylls such as violaxanthin, neoxanthin, and antheraxanthin, along with β-carotene are also present in fruits (Minguez-Mosquera and Hornero-Mendez, 1994b, Hornero-Méndez et al., 2000, Berry et al., 2019). Whilst carotenoid content in harvested pepper fruit has been studied previously (Berry et al., 2019), the change in carotenoid content during post-harvest storage, defined as carotenoid retention, has been understudied to date. Identification of genotypes which retain high levels of carotenoids is crucial to elucidate the mechanisms underlying this trait. The study presented here has demonstrated that pepper genotypes can retain carotenoids to different extents following post-harvest storage. Interestingly, some pepper fruit genotypes showed an increase in carotenoid content during post-harvest storage, including genotypes R5 and R6, which showed carotenoid increases of 48% and 24% respectively, following oven drying. This increase in carotenoids during post-harvest storage has previously been observed in pepper varieties (Park and Lee, 1975, Minguez-Mosquera and Hornero-Mendez, 1994a, Mínguez-Mosquera et al., 2000).

Carotenoid retention is not only a key quality trait in pepper, but also in other crop species. Carotenoid losses during post-harvest storage are well reported in a variety of crop species (Burt et al., 2010, Bechoff et al., 2011), and this affects both the colour quality and nutritional quality. Elevated carotenoid content has been engineered in a variety of crop plants, including rice (Beyer et al., 2002), maize (Zhu et al., 2008), and cassava (Welsch et al., 2010). Whilst these studies have been successful in engineering elevated carotenoid content, and consequently, increased provitamin A capacity, in these crops, significant challenges have been encountered due to the fact that these crops do not retain these carotenoids during post-harvest storage (Hidalgo and Brandolini, 2008, Ortiz et al., 2016).

### The pepper fruit surface structure is associated with the carotenoid retention trait

The cuticle of fruit species has been acknowledged as a modulator of post-harvest quality (Lara et al., 2014), influencing the shelf-life potential of fruit crops (Lara et al., 2019). This study proposes the pepper fruit surface, and specifically the fruit cuticle, as a key structural feature in protecting the fruit against carotenoid losses during post-harvest storage. The carotenoid retention phenotype is associated with the fruit surface structure of pepper, which was noted as genotypes characterised as low carotenoid retention tended to show a ‘wrinkled’ or ‘cracked’ fruit surface upon fruit post-harvest drying, whilst genotypes characterised as high carotenoid retention tended to show a smooth fruit surface upon drying. Ferulic acid has previously been demonstrated to be associated with suberin deposition in melon fruits (Cohen et al., 2019). Suberin deposition occurs in response to wounding, to seal the compromised tissue. Interestingly, high levels of ferulic acid were observed in genotype R4, which further displayed wrinkling upon drying. Consequently, the observed high levels of ferulic acid may be associated with suberin deposition in response to the wrinkling, and hence cracking, of the fruit surface upon drying.

Upon further inspection of the fruit surface using light microscopy, it was noted that pepper genotypes with a smooth surface also possessed a thicker fruit exocarp. The exocarp described here consisted of a lipidic layer in which cells were embedded. The fruit cuticle is normally localised only on the outer surface of epidermal cells (Martin and Rose, 2014), however, in some genotypes studied here, this lipid layer penetrates several cell layers deep within the fruit, and is consequently termed the ‘exocarp’. Microscopic analysis of the tomato cultivar M82 revealed a small amount of sub-epidermal cuticular material deposition (Buda et al., 2009, Yeats et al., 2012), however this did not appear to penetrate multiple cell layers deep into the fruit as presented here in the case of chilli pepper fruits. The cuticular layer has been reported to surround more than a single cell layer below the outermost epidermis in Ailsa Craig tomatoes (Mintz-Oron et al., 2008), however, deposition of cuticular components penetrating below the epidermal cell layer has not been widely reported to the extent demonstrated in this study in pepper fruits. Genotypes possessing a smooth surface tended to have more layers of cells embedded within this lipid layer. A resistance to cuticle cracking has previously been demonstrated to correlate with a thicker fruit cuticle in the case of cherry tomato (Matas et al., 2004).

The cuticle is composed of two components: a cutin-rich section, and embedded cuticular lipophilic components (Yeats and Rose, 2013). Here, increased total cutin monomer content, along with an increase in specific cutin monomers, for example 10,16-dihydroxyhexadecanoic acid, were shown to be correlated with the high carotenoid retention trait in pepper fruits. In contrast, cuticular wax content was not shown to be associated with the carotenoid retention phenotype. This indicates that the thicker exocarp is consisted of a cutin matrix, which may be supported by the finding that the tomato *cd1* mutant, which presents a deficiency in cutin, displayed a significantly thinner cuticular layer with less sub-epidermal deposits than the M82 control (Yeats et al., 2012). These findings suggest that the sub-epidermal deposits observed in the M82 tomato and in the pepper genotypes presented here, are largely comprised of cutin. Cuticular waxes showed no association with exocarp thickness, or with the carotenoid retention trait, and this may be due to their localisation to the outermost layer of the fruit surface. Further, total cuticular waxes have been shown not to correlate with water loss rate in a diverse pepper collection (Parsons et al., 2013). This indicates that cuticle waxes may not play a critical role in traits influenced by the fruit cuticle structure, as demonstrated by their lack of correlation to water loss rate, or to the carotenoid retention trait.

As the cutin matrix penetrates several cell layers deep within the fruit pericarp in the form of sub-epidermal deposits, this suggests that sub-epidermal cell layers, along with epidermal cells, may synthesise cuticular components. Previous identification of attached and detached sub-epidermal globules in the tomato M82 cuticle raised questions regarding the mechanism responsible for depositing cuticular material in sub-epidermal layers (Buda et al., 2009). Two hypotheses were presented: *i)* sub-epidermal cuticular material is derived from epidermal cells and trafficked into sub-epidermal walls; *ii)* sub-epidermal cells, along with epidermal cells, synthesise small amounts of cuticular components (Buda et al., 2009). The data presented here may support the second of these two hypotheses, in that sub-epidermal cells can synthesis cuticular components, as a significant trafficking network would be required in order to explain the depth with which cuticular components penetrate into the fruit pericarp in some of the pepper genotypes described here. Cuticle precursor transport is not entirely understood, however, data presented here suggests that the claim that epidermal cells are responsible for cuticle synthesis and transport (Suh et al., 2005), may need to be reconsidered.

The differential expression of cuticle biosynthesis genes between the high and low carotenoid retention pepper genotypes supports the finding that the fruit surface structure is associated with the carotenoid retention phenotype. Genes identified previously as involved in pepper cuticle biosynthesis (Popovsky-Sarid et al., 2017) were amongst those identified as differentially expressed between the high and low carotenoid retention genotypes. Several genes involved in the biosynthesis of *Arabidopsis thaliana* cuticle (Yeats and Rose, 2013, Mintz-Oron et al., 2008) were found to have orthologs in pepper, which were differentially expressed between the high carotenoid retention genotype, with a thick exocarp, and the low carotenoid retention genotype, with a thinner fruit exocarp. An ortholog of the *Bodyguard* gene (*BDG*), known in *Arabidopsis thaliana* to be involved in cutin biosynthesis (Kurdyukov et al., 2006), is significantly up-regulated in the high retention genotype fruits. This supports the finding that cutin monomer content is also increased, and that the exocarp is thicker, in this high carotenoid retention genotype. However, the *Cutin Deficient 2* (*CD2*) gene, which in tomato has been reported to regulate cutin monomer biosynthesis (Isaacson et al., 2009), is up-regulated in the low carotenoid retention pepper genotype. Despite the fact that a *cd2* tomato mutant was found to decrease in cutin content by 98%, suggesting the crucial role of *CD2* in regulating cutin biosynthesis in tomato fruit (Isaacson et al., 2009), the same influence of this gene may not be exerted in pepper. Differential expression of genes involved in cutin monomer transport, including *ABCG32* (Bessire et al., 2011) and various *LTPs* (Yeats and Rose, 2008) may indicate that transport of these cuticular components is critical to determining the extent of sub-epidermal cutin deposits, and consequently exocarp thickness. Increased expression of these genes appears to result in increased sub-epidermal cutin deposition, and therefore it would seem that cuticle precursor transport is a more critical step in determining cutin content than cutin biosynthesis in pepper fruit. Again, this indicates that previous understanding of epidermal cells being responsible for cuticle precursor synthesis and transport, may need to be reconsidered.

Several genes involved in cuticular wax biosynthesis, including *CER1* and *CER3*, involved in alkane formation from very long chain fatty acid precursors (Bernard et al., 2012), and *MAH1*, a cytochrome P450 with midchain alkane hydroxylase activity (Greer et al., 2007), were also shown to be differentially expressed, although this difference in gene expression does not correlate with cuticular wax content as these waxes were shown not to be associated with the carotenoid retention trait. These contradictory findings between transcript expression and cuticular wax content may be explained by the fact that metabolites may be under pleiotropic control, in which a small number of genes may be responsible for a vast number of metabolites (Chan et al., 2010). Alternatively, as metabolism is a complex and dynamic network in which the products of specific enzymes become substrates for subsequent reactions, steady-state metabolomic analysis lacks detail regarding metabolite flux, and therefore differences in gene expression may not correlate with metabolite content at a single specific time point.

Overexpression of the tomato MIXTA-like MYB transcription factor (*SlMX1*) has previously been shown to result in increased cuticle deposition in the peel, along with an increase in total fruit carotenoid content (Ewas et al., 2016). Whilst homologs of this gene were not found to be differentially expressed between the high and low carotenoid retention pepper genotypes, it suggests the contribution of the cuticle to influencing carotenoid content.

### Implications of the localisation of carotenoids in cells within the fruit exocarp

Interestingly, carotenoids have been shown to be associated with the ripe pepper fruit exocarp. This phenomenon has not previously been observed in pepper, nor in other similar fleshy fruit species, such as tomato. The observation of carotenoids within the lipidic exocarp layer following enzymatic degradation of the cell walls, suggests that the lipid exocarp layer protects carotenoids from degradation. Consequently, the structure of the lipid exocarp directly influences the carotenoid retention phenotype.

Further to this, the ratio of exocarp to whole fruit capsanthin diester content is significantly greater in the high carotenoid retention genotype, R8, compared to the low genotype, R3. This suggests that the waxy, lipophilic environment of the exocarp favours the storage of more non-polar carotenoids, specifically diesters in this case. Esterified carotenoids have been shown to be more stable than their non-esterified counterparts (Schweiggert et al., 2007), and this may explain the increase in total carotenoid content of these cells in the high retention genotype, as esterified carotenoids are less susceptible to oxidative degradation. Whilst the mechanism by which increased esterified carotenoids are accumulated in these exocarp embedded cells, one explanation may be that these cells have an increased capacity for storage of carotenoids in subchromoplast organelles, specifically in fibrillar plastoglobuli, which are well documented in their role in the sequestration and storage of esterified carotenoids in pepper fruit (Deruère et al., 1994). An increase in fibrillar structures would facilitate the accumulation of increased pigment levels within the chromoplasts of high carotenoid retention genotypes, as has previously been demonstrated (Berry et al., 2019).

Carotenoids are localised to cells within exocarp. Spatial localisation of carotenoids within outer fruit layers may influence the visual perception of the fruit, and their localisation within cells embedded within a lipid layer appears to provide protection against degradation.

### Pepper fruit surface protects carotenoids from degradation during post-harvest storage

Treatment of pepper fruits with a cracked surface structure (low carotenoid retention: R3, dried fruits) with hydrogen peroxide as an oxidative agent, resulted in greater degradation of carotenoids than was observed in smooth surface fruits (high carotenoid retention: R8, fresh and dried fruits). Capsorubin, capsanthin, and capsanthin diesters are well characterised as having high quenching capacity for singlet oxygen and hydroxyl free radicals (Nishino et al., 2016). Dried pepper fruits with a cracked cuticle, which showed greater losses in carotenoid content than their smooth cuticle counterparts, following hydrogen peroxide treatment, had greater levels of reactive oxygen species to scavenge. This was presumably due to the cracked cuticle being more permeable to hydrogen peroxide, therefore resulting in increased reactive oxygen species within the fruit responsible for initiating endogenous lipid peroxidation, and consequently increased carotenoid degradation. This provides evidence that the fruit surface structure is critical in protecting the fruit from permeation of reactive oxygen species or their precursors, which initiate lipid peroxidation within the fruit, and consequently carotenoid degradation as carotenoids are scavenged to terminate the lipid peroxidation reactions.

### Biotechnological implications

Significant interest and investment has been made into engineering elevated carotenoid content in crop plants to improve their nutritional properties. Whilst elevated carotenoid content has been successfully engineered (Beyer et al., 2002, Zhu et al., 2008, Welsch et al., 2010), these elevated carotenoid levels tend to be poorly retained throughout post-harvest storage of the crop (Hidalgo and Brandolini, 2008, Ortiz et al., 2016). Analysis of the surface structure of these crops may shed light on their inability to retain carotenoids during post-harvest storage. Previous studies engineering elevated carotenoid crops have focused solely on engineering the carotenoid pathway, whilst insufficient attention has been paid to carotenoid catabolism and changes during post-harvest storage. In addition, because the effects arise under post-harvest treatments they are unlikely to be related to carotenoid derived phytohormones. More recently, studies have considered the role of sequestration and storage mechanisms in engineering elevated carotenoid levels in crop plants (Nogueira et al., 2013). Here, we propose a novel mechanism by which carotenoid content may be controlled in fruit crops. Studies aiming to elevated carotenoid levels in crops should consider the fruit surface structure as a mechanism contributing to carotenoid accumulation and retention.

## Supplementary data

*Table S1*. Pepper diversity panel storage experiment carotenoid amounts.

*Table S2*. Cutin monomer and cuticular wax components of pepper fruit cuticles.

**Fig. S1.**
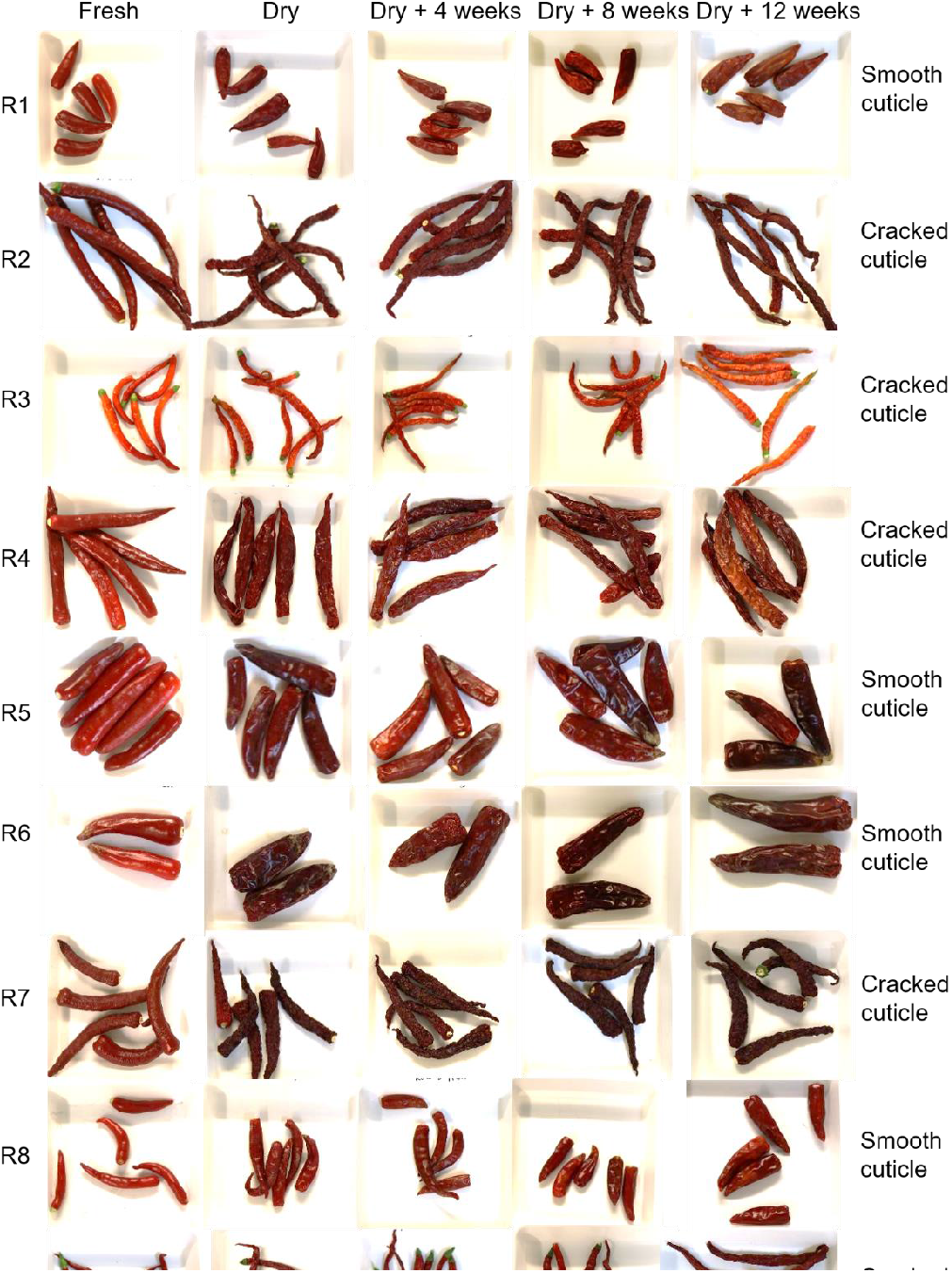
Variation in pepper fruit surface of diversity panel genotypes *appearance following drying and post-harvest storage*.

**Fig. S2.**
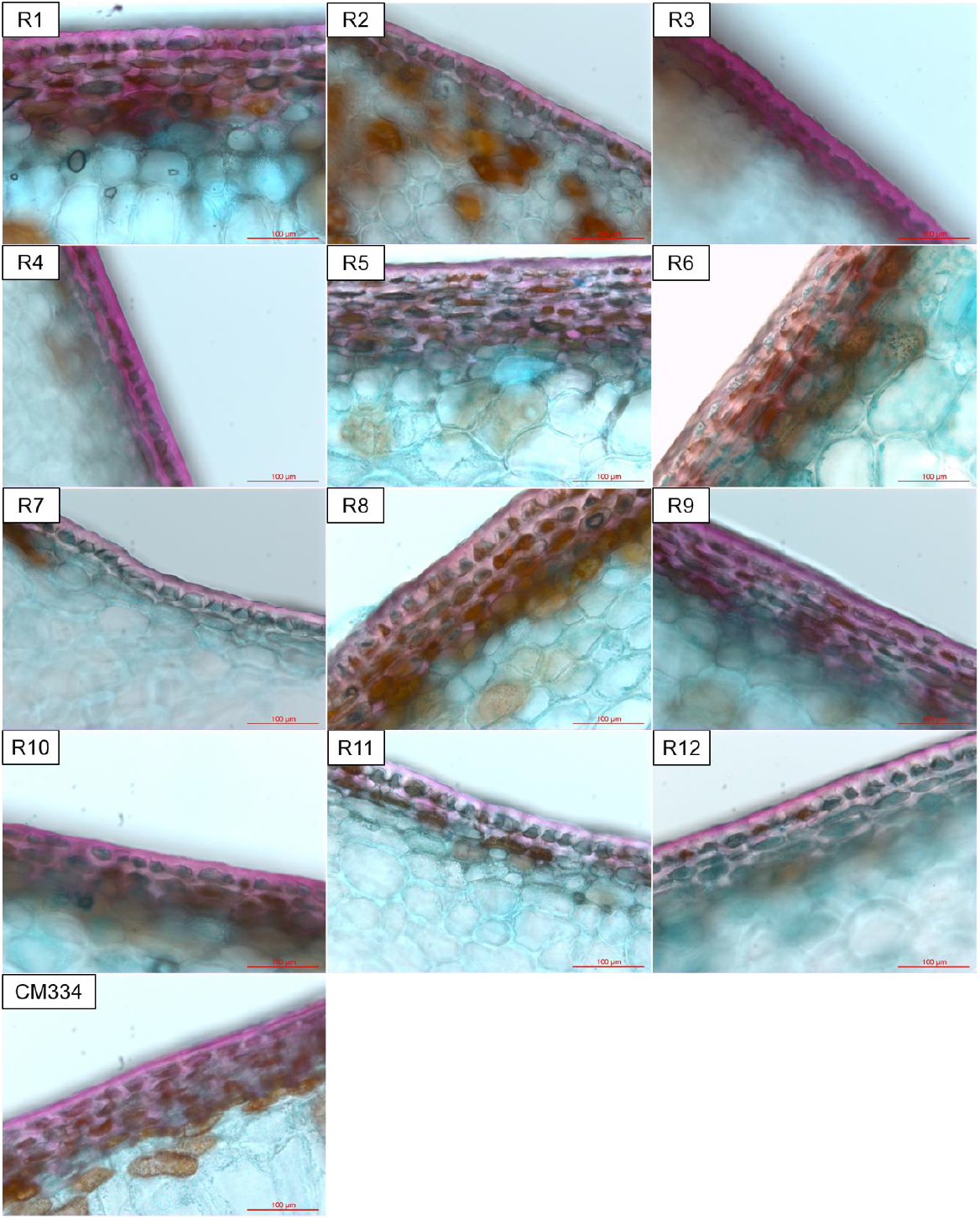
Variation in exocarp structure in pepper diversity panel. Diversity panel, consisting of 13 genotypes, analysed by light microscopy. Exocarp defined as tissue-stained pink using Nile Red stain.

**Fig. S3:**
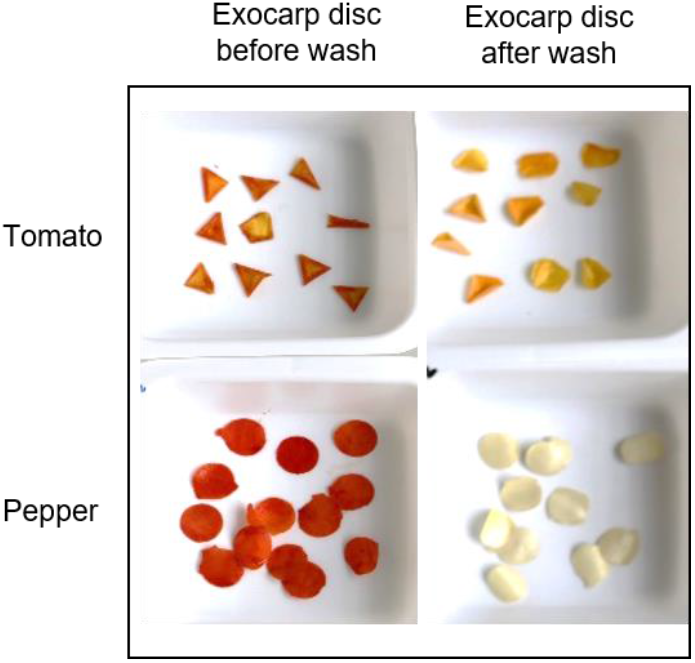
Isolated exocarp discs of pepper and tomato fruits, before and after chloroform washing.

### Experimental Procedures

#### Materials

All chemicals were purchased from Sigma-Aldrich, UK, unless stated otherwise. A commercial panel of 12 chilli pepper genotypes displaying diversity in colour intensity and retention phenotypes was provided by Syngenta. *Capsicum annuum* L. cv. CM334 (Criollo de Morelos 334) was also included in the diversity panel. Material was grown in glasshouses (25 °C, 16/8 h light/dark).

#### Storage conditions

Pepper fruits were harvested when ripe and dried in an oven (30-40 °C) under a 12/12 hour light cycle. Pepper fruits were dried for two weeks before being stored in hessian bags (4 °C) for a period of between 2 – 12 weeks. These conditions were selected in order to replicate the industrial drying and storage process that occurs during commercial growing and storage of chilli peppers.

### Biochemical analyses

#### Carotenoid analysis

Carotenoids were extracted from lyophilised and homogenised chilli pepper powder. 10 mg of tissue was used in the extraction process, using chloroform (500 µL) and HPLC-grade methanol (250 µL). The suspension was incubated on ice, in the dark, before HPLC-grade water (250 µL) was added. A phase separation was created, and the organic layer was collected. Chloroform (500 µL) was again added to the material for extraction, a phase separation was created, and the organic layer was pooled with the initial organic phase. Separation and detection of carotenoids was performed by high performance liquid chromatography with photodiode array detection (HPLC-PDA), using a C30 reverse-phase column (250 × 4.6 mm), purchased from YMC, Wilmington, NC. The solvent system used has been detailed previously (Fraser et al., 2000). Carotenoids isolated from pepper fruit exocarp discs (1 cm) were extracted by washing discs in chloroform and methanol (1:1 ratio; 10 mL) in dark conditions, on a rotator. Wash solution was replaced every 24 hours, and all solvent used for extraction was pooled and evaporated. Exocarp carotenoids were analysed by HPLC-PDA, as described.

#### Cuticle component analysis

*i) Cutin monomer analysis*. 1 cm discs of fruit pericarp tissue were dissected, and cuticle tissue was isolated using pectinase (1.5 % w/v), cellulase (0.1 % w/v), in citrate buffer (0.2 mM, pH 3.7), with sodium azide (1 mM). Samples were incubated, shaking in dark conditions (35 °C; 4 days’ 100 rpm). Cutin monomer extraction was performed as previously described (Cohen et al., 2019). Analysis of extracted cutin monomers was performed using a gas chromatography mass spectrometry (GC-MS) system (Agilent 7683 autosampler, 7890A gas chromatograph, and 5975C mass spectrometer).

*ii) Cuticle wax analysis*. 1 cm fruit discs were dissected and dipped in chloroform (10 mL; 10 s) with cuticle side facing downwards into solvent to extract cuticle bound waxes. Deuterated triacontane (C_30_, 10 µg) was added as an internal standard. 10 discs per biological sample were dipped. Following cuticle wax extraction, the extract was transferred to a cleaned glass vial and dried under nitrogen. Extracts were resuspended in fresh chloroform (500 µL) and transferred to a glass GC-MS vial. Extracts were dried under nitrogen, and derivatised for analysis using pyridine (30 µL) and N-Methyl-N-(trimethylsilyl) trifluoroacetamide (MSTFA; 70 µL). Samples were incubated (40 °C; 1 h) before analysis. Analysis of cuticle waxes was performed using a GC-MS system (Agilent 7820A gas chromatograph, and 5977B mass spectrometer).

For both cutin monomer and cuticle wax analysis, a DB-1HT (Agilent, J+W 122-1131; 30 m x 250 µm x 0.1 µm) was used, with a flow rate of 1.2 mL/minute. Samples were injected in splitless mode at 200 °C, with helium as the carrier gas. The oven temperature was held at 70 °C for 2 minutes, before ramping at 10 °C /minute to 150 °C, and then ramping at 3 °C /minute to 310 °C. The final temperature was held for a further 20 minutes. Mass spectrometry was performed in full scan mode. Identification of chromatogram components was performed by comparing mass spectra to literature reported spectra, in-house libraries, and the NIST 2.0 MS library. Quantification was performed by calculating peak areas relative to the internal standard.

### Fruit microscopy

#### Light microscopy

Fresh fruit were harvested, and hand sectioning was used to dissect cross sections from the middle of the fruit. Sections were collected in distilled water. Nile Red (0.5 mg/mL) in acetone was used to stain lipids, and Fast Green (0.5 % w/v) in ethanol (90 %) was used to stain sub-epidermal cell layers. Sections were washed in distilled water, before mounting on glass slides. A Leica DM 500 compound light microscope was used for visualisation, with 10x and 20x objective lenses. IMS (Imagic, Imaging Ltd.) software was used to capture images.

Exocarp thickness was measured using ImageJ software. Exocarp thickness was measured using light microscopy images. Exocarp was defined as area stained pink following Nile Red staining. Three images per genotype were measured, and six measurements per image were recorded.

#### Cryo-Scanning electron microscopy

Cross sections of fruit were sectioned, frozen in liquid nitrogen, and then sublimed (2 minutes), and coated with platinum. Cryo-scanning electron microscopy (Cryo-SEM) was performed using a Quorum Technologies PP3010T Cryo-SEM system.

#### Scanning electron microscopy

Fruit samples were mounted on aluminium pin stubs (SEM sample supports), using adhesive carbon tabs. Scanning electron microscopy (SEM) was carried out using a Zeiss Evo LS15 scanning electron microscope. Sample imaging was carried out using variable pressure mode, and images were captured via a C2D detector.

#### Transcriptomic analysis

Total RNA was extracted from fruit epidermal cells. Abrasive paper was used to isolate epidermal cells, and RNA was extracted and purified using TRIzol (Invitrogen, UK), and PureLink RNA Minikit (Thermo Fisher Scientific, UK) according to manufacturer’s instructions. RNA was treated with TURBO DNase (ThermoFisher Scientific) to remove contaminating genomic DNA. Following quality control, an mRNA library was prepared using NEBNext Ultra II RNA Library Prep with sample purification beads, and in-house (Syngenta Ltd., Research Triangle Park, USA) 8 base pair indexes. Libraries were sequenced using a Hi-Seq (Illumina). Gene count was normalised using the R package EDASeq v2.16.3 (Risso, 2013), and filtered. Reads were mapped to the published CM334 chilli pepper genome (Kim et al., 2014). The unprocessed transcriptomic data have been deposited with NCBI and can be accessed at www.dataview.ncbi.gov/object/PRJNA640935.

#### Hydrogen peroxide fruit dip

Fruits were harvested and dried as previously described. Fruits were treated with varying concentrations of hydrogen peroxide, either before or after fruit drying, and then stored (4 °C) for one month. Carotenoid content was measured using HPLC-PDA analysis.

#### Data processing and statistical treatment

All experiments typically used three to six biological replicates, unless stated otherwise. Graphs were compiled using GraphPad Prism 8 software, and means, standard deviation, and standard error were calculated using Excel (Microsoft). Significance testing, including Student’s *t* tests and ANOVA were carried out using XLstat software (Addinsoft). Student’s *t* test significances were represented as follows: P < 0.05: *, P < 0.01: **.

## Acknowledgements

The work was supported through a Biotechnology and Biological Sciences Research Council iCASE with Syngenta Ltd., to ACH and PDF; Project BB/P001742/1) and Weizmann-UK, making connections funding. A.A. is the incumbent of the Peter J. Cohn Professorial Chair. Dr Chris Stein (Syngenta, Ltd., UK) is thanked for advice and assistance with microscopy and Dr Julie Green (Syngenta, Ltd., USA), for bioinformatics assistance and expertise. Dr Charles Baxter is thanked for initial project discussions.

## Availability of data and materials

Processed data is available in the manuscript and appendices. The unprocessed data have been deposited with NCBI and can be accessed at www.dataview.ncbi.gov/object/PRJNA640935.

## Author contributions

Conceptualisation (AH, DVR & PDF), methodology (AH, PDF, AA & HC), formal analysis (AH), investigation (AH), resources (PDF & AA), data curation (AH), supervision (PDF, DVR, AA & HC), writing-original draft (AH), Writing-review and editing (AH, HC, DVR, AA, PDF), visualisation (AH), project administration (PDF), and funding acquisition (PDF, DVR & AA).

## Declaration of competing interest

The authors declare that they have no known competing financial interests or personal relationships that could have appeared to influence the work reported in this paper.

## Notes

### Competing Interest Statement

The authors have declared no competing interest.

