## Supplementray material for "Carotenoid retention during post-harvest storage of *Capsicum annuum*: the role of the fruit surface structure"

### Supplementary Materials

**Table S1: Pepper d panel storage experiment carotenoid amounts. Carotenoid amounts of 13 diverse pepper genotypes analysed by HPLC-PDA, at intervals throughout post-harvest storage. Fresh: immediately following harvest; Dry: following post-harvest oven drying; Dry + 4 weeks: following post-harvest drying and 4 weeks cold storage; Dry + 8 weeks: following post-harvest drying and 8 weeks cold storage; Dry + 12 weeks: following post-harvest drying and 12 weeks cold storage. Three biological replicates per sample analysed. Mean ± SEM reported.**

| R1 | Fresh | | Dry | | Dry + 4 weeks | | Dry + 8 weeks | | Dry + 12 weeks | |
|---|---|---|---|---|---|---|---|---|---|---|
| | Mean µg/g | ± SE | Mean µg/g | ± SE | Mean µg/g | ± SE | Mean µg/g | ± SE | Mean µg/g | ± SE |
| Violaxanthin | 133.62 | 5.60 | 102.27 | 3.63 | 87.59 | 5.65 | 94.71 | 5.77 | 98.68 | 10.23 |
| Neoxanthin | 78.95 | 1.70 | 74.79 | 1.76 | 68.00 | 2.14 | 70.09 | 3.01 | 71.85 | 2.01 |
| Antheraxanthin | 51.43 | 4.22 | 41.15 | 2.42 | 31.89 | 2.47 | 38.88 | 3.28 | 35.20 | 3.84 |
| Capsanthin | 558.06 | 32.92 | 392.48 | 4.87 | 359.49 | 14.36 | 380.75 | 14.63 | 374.79 | 18.79 |
| Zeaxanthin | 28.73 | 1.73 | 15.36 | 0.95 | 13.57 | 0.60 | 14.88 | 0.95 | 14.55 | 0.51 |
| Antheraxanthin (monoester) | 179.54 | 9.36 | 206.15 | 17.93 | 169.52 | 26.41 | 196.27 | 21.35 | 160.87 | 36.59 |
| Capsanthin (monoester) | 1059.93 | 89.57 | 1218.41 | 154.24 | 1578.98 | 504.33 | 1404.84 | 122.88 | 1198.25 | 104.13 |
| β-carotene | 182.07 | 41.58 | 150.40 | 25.83 | 76.90 | 6.45 | 102.96 | 11.69 | 121.64 | 31.68 |
| β-cryptoxanthin | 53.37 | 13.32 | 60.92 | 7.98 | 42.13 | 4.45 | 45.47 | 3.71 | 44.59 | 5.15 |
| Capsanthin (diester) | 1921.74 | 102.52 | 2371.96 | 141.13 | 1685.76 | 206.91 | 2399.49 | 169.93 | 2113.95 | 122.39 |
| Zeaxanthin (diester) | 124.88 | 11.28 | 97.15 | 31.73 | 87.64 | 27.92 | 82.11 | 30.11 | 86.66 | 22.18 |
| Capsorubin (diester) | 0.00 | 0.00 | 0.00 | 0.00 | 0.00 | 0.00 | 0.00 | 0.00 | 0.00 | 0.00 |
| Total | 4327.78 | 241.79 | 4710.73 | 303.69 | 4201.46 | 398.33 | 4807.08 | 289.60 | 4288.14 | 169.20 |

| R2 | Fresh | | Dry | | Dry + 4 weeks | | Dry + 8 weeks | | Dry + 12 weeks | |
|---|---|---|---|---|---|---|---|---|---|---|
| | Mean µg/g | ± SE | Mean µg/g | ± SE | Mean µg/g | ± SE | Mean µg/g | ± SE | Mean µg/g | ± SE |
| Violaxanthin | 272.76 | 15.85 | 157.17 | 9.03 | 166.99 | 12.88 | 153.69 | 7.54 | 167.43 | 11.55 |
| Neoxanthin | 138.23 | 9.99 | 86.42 | 0.73 | 90.82 | 5.08 | 89.79 | 3.34 | 95.91 | 6.12 |
| Antheraxanthin | 89.63 | 3.59 | 74.79 | 3.64 | 103.55 | 16.36 | 85.94 | 9.00 | 86.19 | 6.54 |
| Capsanthin | 956.66 | 50.12 | 497.74 | 29.86 | 541.22 | 43.49 | 560.08 | 57.71 | 567.19 | 36.76 |
| Zeaxanthin | 47.01 | 1.35 | 29.38 | 2.96 | 33.34 | 2.67 | 33.01 | 3.78 | 31.25 | 2.62 |
| Antheraxanthin (monoester) | 792.21 | 24.07 | 789.91 | 43.76 | 758.91 | 133.27 | 725.04 | 10.43 | 800.38 | 76.32 |
| Capsanthin (monoester) | 2737.32 | 144.95 | 2919.39 | 121.53 | 3276.51 | 106.10 | 3354.88 | 256.82 | 3166.21 | 397.30 |
| β-carotene | 977.65 | 18.26 | 597.51 | 39.37 | 794.92 | 97.40 | 673.69 | 13.57 | 575.55 | 98.38 |
| β-cryptoxanthin | 207.26 | 71.45 | 156.29 | 20.63 | 217.99 | 60.35 | 175.89 | 23.89 | 187.25 | 33.77 |
| Capsanthin (diester) | 6189.50 | 719.13 | 6652.47 | 414.44 | 6906.83 | 497.12 | 6596.64 | 84.75 | 6858.15 | 534.94 |
| Zeaxanthin (diester) | 305.49 | 50.60 | 233.87 | 8.10 | 334.95 | 13.35 | 375.63 | 76.60 | 342.51 | 136.81 |
| Capsorubin (diester) | 0.00 | 0.00 | 0.00 | 0.00 | 0.00 | 0.00 | 0.00 | 0.00 | 0.00 | 0.00 |
| Total | 12713.72 | 895.11 | 12194.95 | 489.80 | 13226.02 | 888.23 | 12824.26 | 358.10 | 12878.01 | 1244.31 |

| R3 | Fresh | | Dry | | Dry + 4 weeks | | Dry + 8 weeks | | Dry + 12 weeks | |
|---|---|---|---|---|---|---|---|---|---|---|
| | Mean µg/g | ± SE | Mean µg/g | ± SE | Mean µg/g | ± SE | Mean µg/g | ± SE | Mean µg/g | ± SE |
| Violaxanthin | 54.58 | 0.98 | 55.45 | 0.32 | 54.73 | 1.36 | 58.46 | 0.04 | 55.45 | 1.21 |
| Neoxanthin | 0.00 | 0.00 | 0.00 | 0.00 | 0.00 | 0.00 | 52.06 | 0.43 | 0.00 | 0.00 |
| Antheraxanthin | 12.83 | 0.75 | 12.79 | 0.74 | 10.54 | 0.14 | 13.00 | 0.78 | 10.92 | 1.10 |
| Capsanthin | 273.02 | 2.56 | 258.93 | 5.28 | 253.25 | 2.71 | 258.25 | 0.16 | 250.17 | 6.45 |
| Zeaxanthin | 24.87 | 1.63 | 16.53 | 0.96 | 13.72 | 0.67 | 15.32 | 0.83 | 14.33 | 1.81 |
| Antheraxanthin (monoester) | 103.23 | 19.01 | 94.71 | 4.72 | 90.23 | 15.07 | 78.98 | 7.81 | 77.21 | 14.40 |
| Capsanthin (monoester) | 361.72 | 14.19 | 399.76 | 34.62 | 403.86 | 75.73 | 373.35 | 3.07 | 358.99 | 41.01 |
| β-carotene | 173.27 | 25.78 | 156.47 | 8.58 | 130.76 | 30.38 | 98.94 | 15.69 | 128.70 | 26.59 |
| β-cryptoxanthin | 31.37 | 3.36 | 23.22 | 0.32 | 25.55 | 3.23 | 24.22 | 0.68 | 26.75 | 0.00 |
| Capsanthin (diester) | 1276.21 | 43.94 | 1248.55 | 25.78 | 1104.41 | 88.27 | 1123.46 | 2.33 | 1034.16 | 186.13 |
| Zeaxanthin (diester) | 179.46 | 51.30 | 133.79 | 10.49 | 129.32 | 22.43 | 166.66 | 51.82 | 150.05 | 31.62 |
| Capsorubin (diester) | 252.77 | 0.00 | 0.00 | 0.00 | 0.00 | 0.00 | 0.00 | 0.00 | 0.00 | 0.00 |
| Total | 2574.82 | 77.98 | 2392.46 | 12.06 | 2189.62 | 232.94 | 2262.70 | 42.26 | 2088.89 | 313.35 |

| R4 | Fresh | | Dry | | Dry + 4 weeks | | Dry + 8 weeks | | Dry + 12 weeks | |
|---|---|---|---|---|---|---|---|---|---|---|
| | Mean µg/g | ± SE | Mean µg/g | ± SE | Mean µg/g | ± SE | Mean µg/g | ± SE | Mean µg/g | ± SE |
| Violaxanthin | 178.41 | 16.20 | 156.78 | 12.04 | 151.13 | 1.00 | 135.91 | 6.32 | 126.75 | 25.10 |
| Neoxanthin | 80.03 | 7.17 | 75.46 | 4.89 | 77.26 | 1.24 | 74.47 | 4.16 | 68.00 | 5.39 |
| Antheraxanthin | 77.18 | 9.04 | 71.35 | 7.07 | 72.98 | 2.59 | 65.65 | 6.05 | 58.77 | 19.91 |
| Capsanthin | 431.64 | 34.62 | 346.20 | 22.31 | 352.50 | 15.29 | 349.95 | 19.07 | 323.00 | 39.49 |
| Zeaxanthin | 39.59 | 4.18 | 24.64 | 1.82 | 24.09 | 1.40 | 23.36 | 1.53 | 19.81 | 4.36 |

| | | | | | | | | | | |
|---|---|---|---|---|---|---|---|---|---|---|
| Antheraxanthin (monoester) | 500.44 | 71.94 | 290.65 | 19.73 | 318.55 | 37.17 | 305.89 | 36.93 | 220.06 | 8.40 |
| Capsanthin (monoester) | 1966.04 | 454.75 | 2415.61 | 230.73 | 2253.13 | 240.08 | 2210.96 | 198.25 | 1758.08 | 456.16 |
| β-carotene | 243.99 | 63.53 | 144.37 | 13.58 | 271.61 | 42.48 | 93.75 | 22.32 | 144.46 | 16.98 |
| β-cryptoxanthin | 86.48 | 4.59 | 117.54 | 8.26 | 102.20 | 22.55 | 71.35 | 24.74 | 95.57 | 17.29 |
| Capsanthin (diester) | 4103.99 | 543.71 | 4505.22 | 280.24 | 4608.65 | 439.72 | 3662.00 | 144.78 | 3310.66 | 533.99 |
| Zeaxanthin (diester) | 194.57 | 32.56 | 73.34 | 24.65 | 137.63 | 57.30 | 76.95 | 23.60 | 75.69 | 21.69 |
| Capsorubin (diester) | 0.00 | 0.00 | 0.00 | 0.00 | 0.00 | 0.00 | 317.65 | 22.55 | 0.00 | 0.00 |
| Total | 7902.35 | 964.11 | 8221.15 | 567.16 | 8343.97 | 537.49 | 7282.00 | 167.49 | 6200.86 | 1120.50 |

| **R5** | **Fresh** | | **Dry** | | **Dry + 4 weeks** | | **Dry + 8 weeks** | | **Dry + 12 weeks** | |
|---|---|---|---|---|---|---|---|---|---|---|
| | **Mean µg/g** | **± SE** | **Mean µg/g** | **± SE** | **Mean µg/g** | **± SE** | **Mean µg/g** | **± SE** | **Mean µg/g** | **± SE** |
| Violaxanthin | 94.42 | 8.73 | 96.61 | 4.77 | 97.60 | 1.84 | 92.88 | 3.62 | 96.01 | 6.23 |
| Neoxanthin | 61.04 | 1.19 | 68.78 | 1.13 | 71.95 | 6.63 | 69.77 | 5.13 | 65.50 | 2.39 |
| Antheraxanthin | 27.14 | 3.54 | 44.77 | 5.24 | 51.27 | 13.66 | 56.90 | 15.35 | 37.56 | 3.09 |
| Capsanthin | 365.90 | 28.46 | 363.27 | 24.13 | 365.67 | 37.73 | 386.70 | 44.22 | 369.39 | 18.62 |
| Zeaxanthin | 19.94 | 2.71 | 14.57 | 0.98 | 13.14 | 0.19 | 13.67 | 1.39 | 14.31 | 1.17 |
| Antheraxanthin (monoester) | 203.66 | 15.87 | 287.01 | 40.95 | 192.20 | 41.19 | 190.36 | 27.97 | 266.56 | 24.17 |
| Capsanthin (monoester) | 586.31 | 41.96 | 1596.09 | 56.45 | 1139.92 | 42.29 | 1186.38 | 164.02 | 1198.06 | 183.07 |
| β-carotene | 197.55 | 15.91 | 91.58 | 3.45 | 218.16 | 81.31 | 161.84 | 33.70 | 223.07 | 62.32 |
| β-cryptoxanthin | 33.05 | 0.00 | 28.83 | 0.83 | 41.51 | 10.18 | 47.97 | 4.30 | 50.77 | 0.67 |
| Capsanthin (diester) | 2203.65 | 107.82 | 3305.36 | 145.06 | 2657.02 | 550.72 | 2355.14 | 348.71 | 3155.47 | 284.90 |
| Zeaxanthin (diester) | 73.07 | 7.42 | 93.34 | 17.85 | 68.15 | 18.60 | 69.40 | 27.16 | 140.85 | 34.99 |
| Capsorubin (diester) | 0.00 | 0.00 | 0.00 | 0.00 | 238.86 | 0.00 | 255.38 | 0.00 | 0.00 | 0.00 |
| Total | 3812.21 | 223.04 | 5935.08 | 196.74 | 4963.68 | 624.80 | 4685.17 | 272.96 | 5585.53 | 494.29 |

| **R6** | **Fresh** | | **Dry** | | **Dry + 4 weeks** | | **Dry + 8 weeks** | | **Dry + 12 weeks** | |
|---|---|---|---|---|---|---|---|---|---|---|
| | **Mean µg/g** | **± SE** | **Mean µg/g** | **± SE** | **Mean µg/g** | **± SE** | **Mean µg/g** | **± SE** | **Mean µg/g** | **± SE** |
| Violaxanthin | 91.82 | 7.96 | 94.02 | 7.03 | 87.90 | 2.35 | 111.96 | 5.86 | 106.20 | 11.12 |
| Neoxanthin | 60.96 | 1.96 | 68.42 | 2.79 | 64.35 | 2.41 | 70.15 | 0.46 | 70.17 | 2.93 |
| Antheraxanthin | 24.23 | 0.98 | 25.56 | 1.89 | 23.62 | 1.26 | 29.31 | 4.34 | 26.14 | 1.85 |
| Capsanthin | 348.80 | 6.77 | 305.31 | 13.14 | 303.61 | 9.66 | 316.70 | 5.94 | 317.74 | 6.61 |
| Zeaxanthin | 22.78 | 1.53 | 15.85 | 0.77 | 17.20 | 3.38 | 16.71 | 1.46 | 16.29 | 1.01 |
| Antheraxanthin (monoester) | 342.61 | 11.46 | 418.60 | 45.70 | 421.02 | 16.17 | 465.86 | 66.00 | 366.60 | 55.48 |
| Capsanthin (monoester) | 748.13 | 40.29 | 1208.91 | 155.00 | 1183.41 | 148.28 | 1258.67 | 28.90 | 1240.50 | 106.03 |
| β-carotene | 284.69 | 20.91 | 288.33 | 33.17 | 300.75 | 59.02 | 391.35 | 56.26 | 326.07 | 64.29 |
| β-cryptoxanthin | 62.66 | 20.54 | 87.44 | 10.71 | 51.18 | 7.89 | 111.58 | 15.15 | 110.81 | 17.16 |
| Capsanthin (diester) | 3525.77 | 84.91 | 4867.55 | 257.18 | 4389.55 | 213.54 | 4903.55 | 546.65 | 4359.45 | 493.68 |
| Zeaxanthin (diester) | 217.25 | 16.25 | 217.78 | 28.31 | 202.20 | 63.94 | 232.41 | 30.53 | 160.23 | 32.35 |
| Capsorubin (diester) | 0.00 | 0.00 | 0.00 | 0.00 | 262.10 | 0.00 | 0.00 | 0.00 | 0.00 | 0.00 |
| Total | 5699.11 | 98.50 | 7543.61 | 370.74 | 7081.41 | 354.89 | 7870.92 | 737.82 | 7064.79 | 706.14 |

| **R7** | **Fresh** | | **Dry** | | **Dry + 4 weeks** | | **Dry + 8 weeks** | | **Dry + 12 weeks** | |
|---|---|---|---|---|---|---|---|---|---|---|
| | **Mean µg/g** | **± SE** | **Mean µg/g** | **± SE** | **Mean µg/g** | **± SE** | **Mean µg/g** | **± SE** | **Mean µg/g** | **± SE** |
| Violaxanthin | 361.54 | 29.77 | 180.33 | 34.82 | 181.22 | 9.42 | 261.74 | 27.97 | 202.30 | 44.52 |
| Neoxanthin | 134.71 | 10.04 | 94.31 | 8.82 | 94.92 | 2.46 | 108.43 | 13.38 | 97.04 | 7.19 |
| Antheraxanthin | 125.03 | 7.35 | 87.74 | 9.31 | 91.02 | 1.99 | 110.57 | 16.78 | 93.53 | 6.62 |
| Capsanthin | 722.36 | 23.81 | 510.53 | 41.43 | 537.08 | 11.91 | 567.88 | 37.60 | 495.22 | 15.83 |
| Zeaxanthin | 81.89 | 13.41 | 43.11 | 2.41 | 37.08 | 1.00 | 39.12 | 0.93 | 33.94 | 3.79 |
| Antheraxanthin (monoester) | 1077.27 | 105.06 | 642.30 | 131.90 | 888.64 | 62.55 | 1105.17 | 89.53 | 783.69 | 184.55 |
| Capsanthin (monoester) | 4147.52 | 195.76 | 3542.08 | 265.51 | 3265.66 | 202.81 | 3993.00 | 581.26 | 3586.68 | 202.16 |
| β-carotene | 1690.45 | 74.28 | 860.49 | 55.68 | 838.74 | 126.29 | 908.75 | 111.99 | 722.22 | 108.63 |
| β-cryptoxanthin | 425.33 | 67.68 | 186.05 | 57.10 | 110.19 | 27.89 | 187.99 | 28.22 | 176.04 | 9.45 |
| Capsanthin (diester) | 9028.95 | 346.52 | 6433.29 | 951.51 | 6829.21 | 445.33 | 8333.79 | 1209.85 | 6855.18 | 1100.40 |
| Zeaxanthin (diester) | 448.69 | 23.23 | 431.04 | 42.05 | 400.74 | 110.65 | 446.42 | 46.55 | 249.41 | 50.75 |
| Capsorubin (diester) | 0.00 | 0.00 | 0.00 | 0.00 | 0.00 | 0.00 | 0.00 | 0.00 | 289.91 | 0.00 |
| Total | 18123.22 | 693.16 | 12951.16 | 1507.75 | 13214.09 | 915.17 | 15975.60 | 2170.30 | 13324.45 | 1525.93 |

| **R8** | **Fresh** | | **Dry** | | **Dry + 4 weeks** | | **Dry + 8 weeks** | | **Dry + 12 weeks** | |
|---|---|---|---|---|---|---|---|---|---|---|
| | **Mean µg/g** | **± SE** | **Mean µg/g** | **± SE** | **Mean µg/g** | **± SE** | **Mean µg/g** | **± SE** | **Mean µg/g** | **± SE** |
| Violaxanthin | 82.93 | 6.84 | 78.45 | 2.42 | 75.90 | 1.32 | 74.24 | 7.36 | 79.64 | 5.70 |
| Neoxanthin | 57.05 | 2.59 | 62.01 | 1.90 | 59.93 | 1.70 | 59.08 | 2.28 | 59.81 | 1.73 |

| | | | | | | | | | | |
|---|---|---|---|---|---|---|---|---|---|---|
| Antheraxanthin | 16.74 | 1.37 | 40.29 | 15.49 | 29.34 | 6.31 | 34.34 | 6.65 | 34.52 | 8.59 |
| Capsanthin | 273.52 | 6.31 | 324.93 | 49.15 | 296.28 | 33.39 | 301.55 | 35.26 | 292.07 | 21.07 |
| Zeaxanthin | 13.24 | 1.00 | 13.51 | 2.16 | 12.72 | 0.06 | 11.74 | 1.60 | 11.33 | 1.06 |
| Antheraxanthin (monoester) | 173.99 | 17.37 | 203.38 | 8.56 | 144.18 | 12.51 | 137.85 | 13.10 | 166.24 | 17.59 |
| Capsanthin (monoester) | 517.97 | 92.09 | 897.49 | 198.70 | 849.66 | 141.08 | 864.04 | 182.11 | 701.04 | 96.86 |
| β-carotene | 198.93 | 30.96 | 185.21 | 39.06 | 192.07 | 28.89 | 145.82 | 47.51 | 169.95 | 19.93 |
| β-cryptoxanthin | 37.37 | 8.88 | 65.73 | 11.20 | 41.86 | 6.61 | 61.48 | 27.78 | 54.38 | 12.87 |
| Capsanthin (diester) | 2572.98 | 348.42 | 2776.54 | 15.39 | 2646.71 | 164.73 | 2065.59 | 116.89 | 2254.77 | 174.15 |
| Zeaxanthin (diester) | 144.13 | 21.53 | 100.80 | 27.14 | 86.19 | 28.75 | 82.89 | 15.00 | 117.20 | 33.24 |
| Capsorubin (diester) | 0.00 | 0.00 | 0.00 | 0.00 | 270.72 | 0.00 | 275.34 | 11.51 | 290.02 | 0.00 |
| Total | 4042.19 | 465.70 | 4701.52 | 215.46 | 4475.56 | 358.72 | 3997.42 | 299.41 | 3991.15 | 348.58 |

| **R9** | **Fresh** | | **Dry** | | **Dry + 4 weeks** | | **Dry + 8 weeks** | | **Dry + 12 weeks** | |
|---|---|---|---|---|---|---|---|---|---|---|
| | **Mean µg/g** | **± SE** | **Mean µg/g** | **± SE** | **Mean µg/g** | **± SE** | **Mean µg/g** | **± SE** | **Mean µg/g** | **± SE** |
| Violaxanthin | 175.52 | 18.33 | 113.66 | 10.76 | 111.07 | 2.93 | 105.43 | 0.22 | 110.25 | 10.71 |
| Neoxanthin | 88.34 | 4.54 | 71.02 | 0.34 | 68.72 | 1.72 | 68.07 | 1.27 | 66.46 | 0.66 |
| Antheraxanthin | 53.11 | 4.63 | 36.12 | 1.36 | 31.74 | 2.51 | 29.69 | 1.35 | 32.38 | 4.44 |
| Capsanthin | 501.24 | 59.84 | 302.36 | 2.73 | 296.18 | 12.07 | 297.46 | 6.43 | 302.75 | 9.69 |
| Zeaxanthin | 17.64 | 2.10 | 12.00 | 0.41 | 9.88 | 0.26 | 15.10 | 0.00 | 15.78 | 0.72 |
| Antheraxanthin (monoester) | 373.62 | 39.22 | 358.54 | 30.24 | 291.74 | 4.74 | 260.10 | 11.92 | 299.11 | 73.54 |
| Capsanthin (monoester) | 1975.01 | 215.12 | 1569.54 | 117.61 | 1601.07 | 44.85 | 1281.08 | 74.20 | 1591.28 | 173.15 |
| β-carotene | 112.73 | 16.05 | 116.72 | 18.40 | 68.77 | 12.40 | 35.47 | 5.23 | 46.83 | 12.06 |
| β-cryptoxanthin | 102.16 | 20.52 | 76.22 | 7.29 | 100.79 | 15.57 | 58.45 | 0.88 | 66.20 | 3.78 |
| Capsanthin (diester) | 4043.99 | 490.34 | 3916.26 | 310.13 | 3739.25 | 381.06 | 3007.86 | 76.61 | 3652.01 | 344.14 |
| Zeaxanthin (diester) | 132.47 | 44.22 | 45.75 | 27.08 | 150.35 | 16.90 | 81.89 | 23.12 | 86.54 | 32.08 |
| Capsorubin (diester) | 0.00 | 0.00 | 0.00 | 0.00 | 0.00 | 0.00 | 0.00 | 0.00 | 0.00 | 0.00 |
| Total | 7517.33 | 719.98 | 6552.65 | 421.38 | 6429.26 | 419.21 | 5195.41 | 196.45 | 6205.42 | 593.81 |

| **R10** | **Fresh** | | **Dry** | | **Dry + 4 weeks** | | **Dry + 8 weeks** | | **Dry + 12 weeks** | |
|---|---|---|---|---|---|---|---|---|---|---|
| | **Mean µg/g** | **± SE** | **Mean µg/g** | **± SE** | **Mean µg/g** | **± SE** | **Mean µg/g** | **± SE** | **Mean µg/g** | **± SE** |
| Violaxanthin | 77.87 | 17.71 | 80.86 | 5.87 | 72.36 | 0.60 | 70.54 | 5.05 | 71.08 | 6.23 |
| Neoxanthin | 58.47 | 2.44 | 0.00 | 0.00 | 57.06 | 0.00 | 0.00 | 0.00 | 0.00 | 0.00 |
| Antheraxanthin | 23.29 | 3.61 | 27.59 | 1.53 | 22.81 | 0.76 | 24.16 | 2.63 | 21.43 | 1.84 |
| Capsanthin | 317.94 | 15.71 | 267.50 | 4.12 | 259.40 | 3.80 | 257.91 | 4.78 | 252.69 | 4.61 |
| Zeaxanthin | 22.87 | 0.74 | 14.61 | 0.36 | 13.03 | 0.74 | 14.08 | 1.31 | 13.29 | 0.75 |
| Antheraxanthin (monoester) | 178.47 | 26.11 | 292.80 | 26.04 | 208.76 | 27.62 | 241.83 | 51.40 | 199.30 | 16.92 |
| Capsanthin (monoester) | 557.55 | 81.32 | 958.11 | 73.80 | 778.71 | 38.27 | 866.05 | 99.61 | 755.70 | 76.24 |
| β-carotene | 156.31 | 21.17 | 218.23 | 112.18 | 113.96 | 16.51 | 79.06 | 31.59 | 90.61 | 10.85 |
| β-cryptoxanthin | 51.53 | 15.89 | 37.13 | 3.94 | 83.64 | 12.82 | 34.89 | 2.94 | 56.33 | 4.59 |
| Capsanthin (diester) | 2173.13 | 438.99 | 2821.40 | 173.72 | 2414.16 | 98.23 | 2359.44 | 536.22 | 2330.60 | 60.71 |
| Zeaxanthin (diester) | 250.92 | 84.92 | 119.88 | 33.03 | 131.70 | 18.39 | 110.22 | 31.57 | 191.13 | 73.99 |
| Capsorubin (diester) | 0.00 | 0.00 | 0.00 | 0.00 | 0.00 | 0.00 | 601.46 | 12.76 | 0.00 | 0.00 |
| Total | 3842.39 | 610.55 | 4722.00 | 296.06 | 4093.44 | 135.36 | 4435.65 | 420.36 | 3958.48 | 21.47 |

| **R11** | **Fresh** | | **Dry** | | **Dry + 4 weeks** | | **Dry + 8 weeks** | | **Dry + 12 weeks** | |
|---|---|---|---|---|---|---|---|---|---|---|
| | **Mean µg/g** | **± SE** | **Mean µg/g** | **± SE** | **Mean µg/g** | **± SE** | **Mean µg/g** | **± SE** | **Mean µg/g** | **± SE** |
| Violaxanthin | 109.52 | 12.06 | 104.19 | 7.71 | 93.77 | 8.09 | 90.10 | 0.00 | 113.97 | 5.40 |
| Neoxanthin | 67.46 | 4.12 | 0.00 | 0.00 | 61.62 | 1.38 | 59.11 | 2.33 | 0.00 | 0.00 |
| Antheraxanthin | 38.39 | 2.02 | 39.79 | 5.17 | 35.57 | 6.62 | 36.11 | 0.84 | 39.56 | 3.31 |
| Capsanthin | 322.46 | 12.32 | 308.31 | 12.98 | 294.60 | 5.47 | 297.86 | 9.28 | 307.62 | 5.01 |
| Zeaxanthin | 43.11 | 3.98 | 26.52 | 0.17 | 21.11 | 2.21 | 22.91 | 1.98 | 24.52 | 2.13 |
| Antheraxanthin (monoester) | 360.43 | 20.12 | 214.62 | 29.23 | 223.03 | 17.57 | 170.32 | 74.52 | 172.37 | 29.90 |
| Capsanthin (monoester) | 920.09 | 104.46 | 1383.50 | 122.61 | 1156.58 | 96.76 | 924.61 | 80.15 | 1347.54 | 185.98 |
| β-carotene | 232.02 | 14.50 | 59.99 | 11.09 | 101.89 | 22.66 | 130.20 | 18.61 | 124.38 | 30.01 |
| β-cryptoxanthin | 60.02 | 18.30 | 49.55 | 5.29 | 37.02 | 8.22 | 42.38 | 3.98 | 71.55 | 12.40 |
| Capsanthin (diester) | 3103.48 | 240.56 | 3094.65 | 227.62 | 2641.32 | 312.95 | 2207.83 | 290.45 | 3345.66 | 231.87 |
| Zeaxanthin (diester) | 235.94 | 50.22 | 87.92 | 12.46 | 78.20 | 11.92 | 74.70 | 12.50 | 87.77 | 8.72 |
| Capsorubin (diester) | 0.00 | 0.00 | 239.99 | 0.00 | 0.00 | 0.00 | 252.45 | 8.39 | 261.39 | 0.00 |
| Total | 5436.39 | 378.47 | 5414.32 | 333.91 | 4713.44 | 428.32 | 4263.54 | 399.34 | 5684.06 | 372.93 |

| **R12** | **Fresh** | | **Dry** | | **Dry + 4 weeks** | | **Dry + 8 weeks** | | **Dry + 12 weeks** |
|---|---|---|---|---|---|---|---|---|---|

| | Mean µg/g | ± SE | Mean µg/g | ± SE | Mean µg/g | ± SE | Mean µg/g | ± SE | Mean µg/g | ± SE |
|---|---|---|---|---|---|---|---|---|---|---|
| Violaxanthin | 84.63 | 2.96 | 67.64 | 3.44 | 73.73 | 0.37 | 71.68 | 0.00 | 72.35 | 0.00 |
| Neoxanthin | 60.56 | 1.65 | 0.00 | 0.00 | 0.00 | 0.00 | 54.70 | 0.86 | 0.00 | 0.00 |
| Antheraxanthin | 48.79 | 8.52 | 32.17 | 6.42 | 29.49 | 3.48 | 27.65 | 1.00 | 29.74 | 2.02 |
| Capsanthin | 498.56 | 55.27 | 304.86 | 17.62 | 293.66 | 9.33 | 290.26 | 2.89 | 306.45 | 4.23 |
| Zeaxanthin | 45.01 | 5.01 | 17.31 | 1.29 | 17.53 | 1.83 | 15.01 | 0.28 | 18.56 | 0.44 |
| Antheraxanthin (monoester) | 157.08 | 20.11 | 101.68 | 14.64 | 123.29 | 16.03 | 104.42 | 18.73 | 126.93 | 3.84 |
| Capsanthin (monoester) | 919.52 | 11.41 | 863.21 | 17.25 | 782.92 | 23.46 | 761.55 | 123.54 | 1084.78 | 80.92 |
| β-carotene | 120.87 | 10.09 | 54.69 | 6.61 | 127.22 | 26.95 | 99.69 | 15.97 | 74.78 | 10.22 |
| β-cryptoxanthin | 31.86 | 3.63 | 25.44 | 0.00 | 41.53 | 9.95 | 62.62 | 7.77 | 65.18 | 0.00 |
| Capsanthin (diester) | 1622.87 | 76.81 | 1220.91 | 53.05 | 1622.27 | 119.61 | 1405.11 | 253.40 | 1955.26 | 20.84 |
| Zeaxanthin (diester) | 195.77 | 71.24 | 151.05 | 61.62 | 72.88 | 3.99 | 54.08 | 16.72 | 45.21 | 16.52 |
| Capsorubin (diester) | 0.00 | 0.00 | 0.00 | 0.00 | 0.00 | 0.00 | 0.00 | 0.00 | 0.00 | 0.00 |
| Total | 3757.30 | 71.02 | 2799.45 | 66.95 | 3159.94 | 197.46 | 2910.94 | 444.51 | 3710.48 | 3.41 |

| CM334 | Fresh | | Dry | | Dry + 4 weeks | | Dry + 8 weeks | | Dry + 12 weeks | |
|---|---|---|---|---|---|---|---|---|---|---|
| | Mean µg/g | ± SE | Mean µg/g | ± SE | Mean µg/g | ± SE | Mean µg/g | ± SE | Mean µg/g | ± SE |
| Violaxanthin | 85.69 | 10.77 | 85.55 | 3.16 | 87.11 | 5.28 | 94.01 | 4.07 | 87.09 | 9.38 |
| Neoxanthin | 62.54 | 5.80 | 61.34 | 0.68 | 59.53 | 2.14 | 62.27 | 1.19 | 60.68 | 2.16 |
| Antheraxanthin | 41.46 | 1.52 | 29.78 | 2.21 | 33.95 | 6.51 | 31.13 | 1.73 | 46.11 | 8.94 |
| Capsanthin | 408.76 | 13.68 | 327.06 | 4.34 | 305.24 | 15.21 | 328.17 | 4.44 | 335.17 | 28.94 |
| Zeaxanthin | 43.14 | 9.18 | 12.19 | 0.72 | 11.94 | 0.88 | 11.88 | 0.25 | 13.10 | 1.39 |
| Antheraxanthin (monoester) | 374.87 | 74.55 | 140.07 | 15.17 | 136.86 | 5.77 | 194.80 | 29.14 | 135.42 | 22.72 |
| Capsanthin (monoester) | 1289.40 | 160.75 | 1319.31 | 131.29 | 1194.33 | 132.12 | 1412.55 | 97.41 | 1277.67 | 155.91 |
| β-carotene | 267.04 | 116.33 | 59.66 | 6.85 | 64.05 | 8.31 | 44.44 | 3.26 | 83.82 | 12.33 |
| β-cryptoxanthin | 32.75 | 0.00 | 123.72 | 46.81 | 106.25 | 28.22 | 91.51 | 21.94 | 82.72 | 7.34 |
| Capsanthin (diester) | 3100.26 | 424.85 | 2681.21 | 228.27 | 2496.36 | 166.10 | 2739.53 | 230.89 | 2483.25 | 163.94 |
| Zeaxanthin (diester) | 192.31 | 46.38 | 57.68 | 5.37 | 69.10 | 24.58 | 108.46 | 66.86 | 80.43 | 11.80 |
| Capsorubin (diester) | 0.00 | 0.00 | 0.00 | 0.00 | 0.00 | 0.00 | 312.41 | 26.84 | 0.00 | 0.00 |
| Total | 5826.99 | 665.82 | 4864.99 | 391.56 | 4535.69 | 353.01 | 5395.86 | 354.40 | 4656.41 | 281.09 |

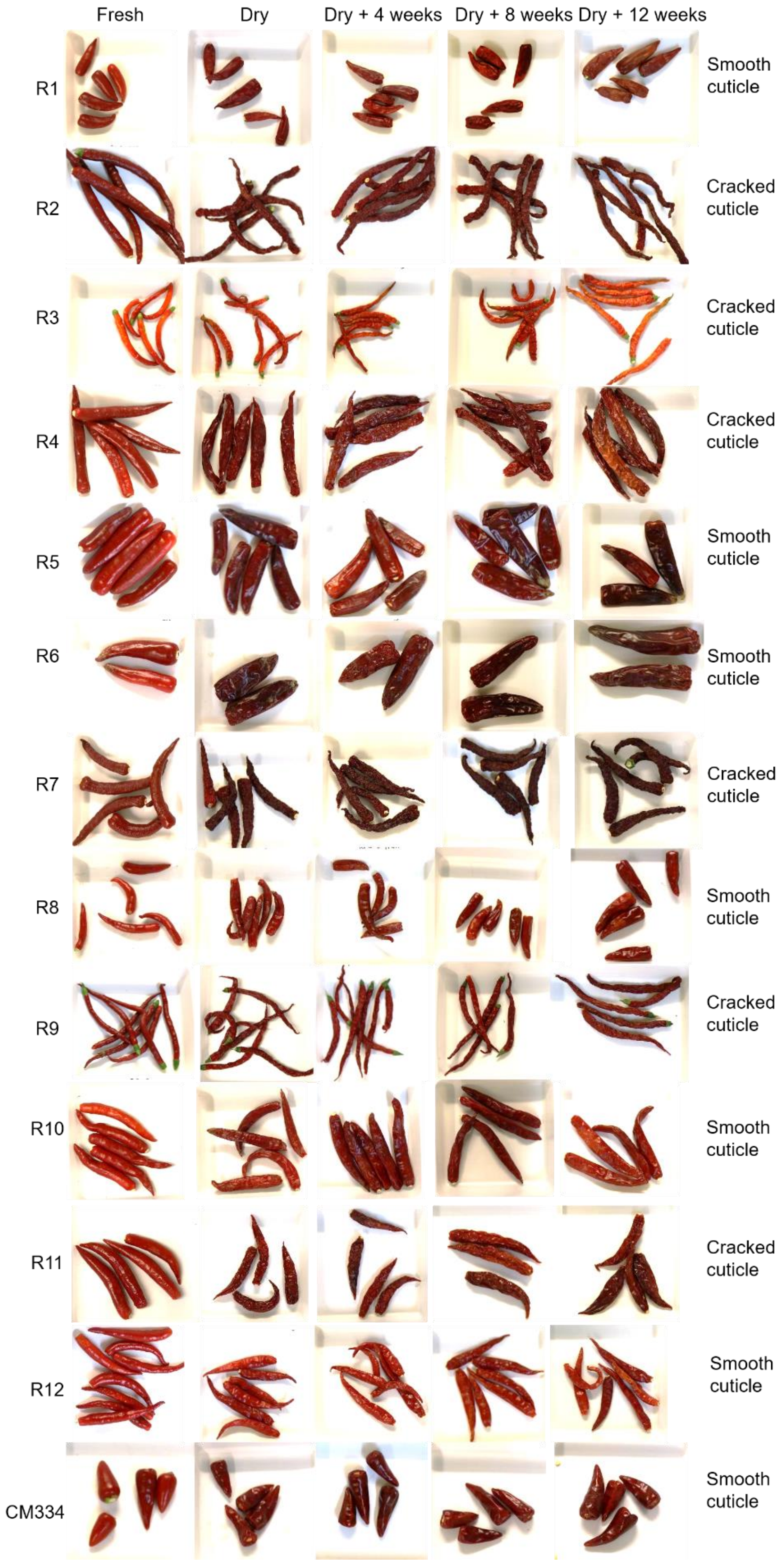

**Fig. S1: Variation in pepper fruit surface of diversity panel genotypes appearance following drying and post-harvest storage.** Fresh: immediately following harvest; Dry: following post-harvest oven drying; Dry + 4 weeks: following post-harvest drying and 4 weeks cold storage; Dry + 8 weeks: following post-harvest drying and 8 weeks cold storage; Dry + 12 weeks: following post-harvest drying and 12 weeks cold storage.

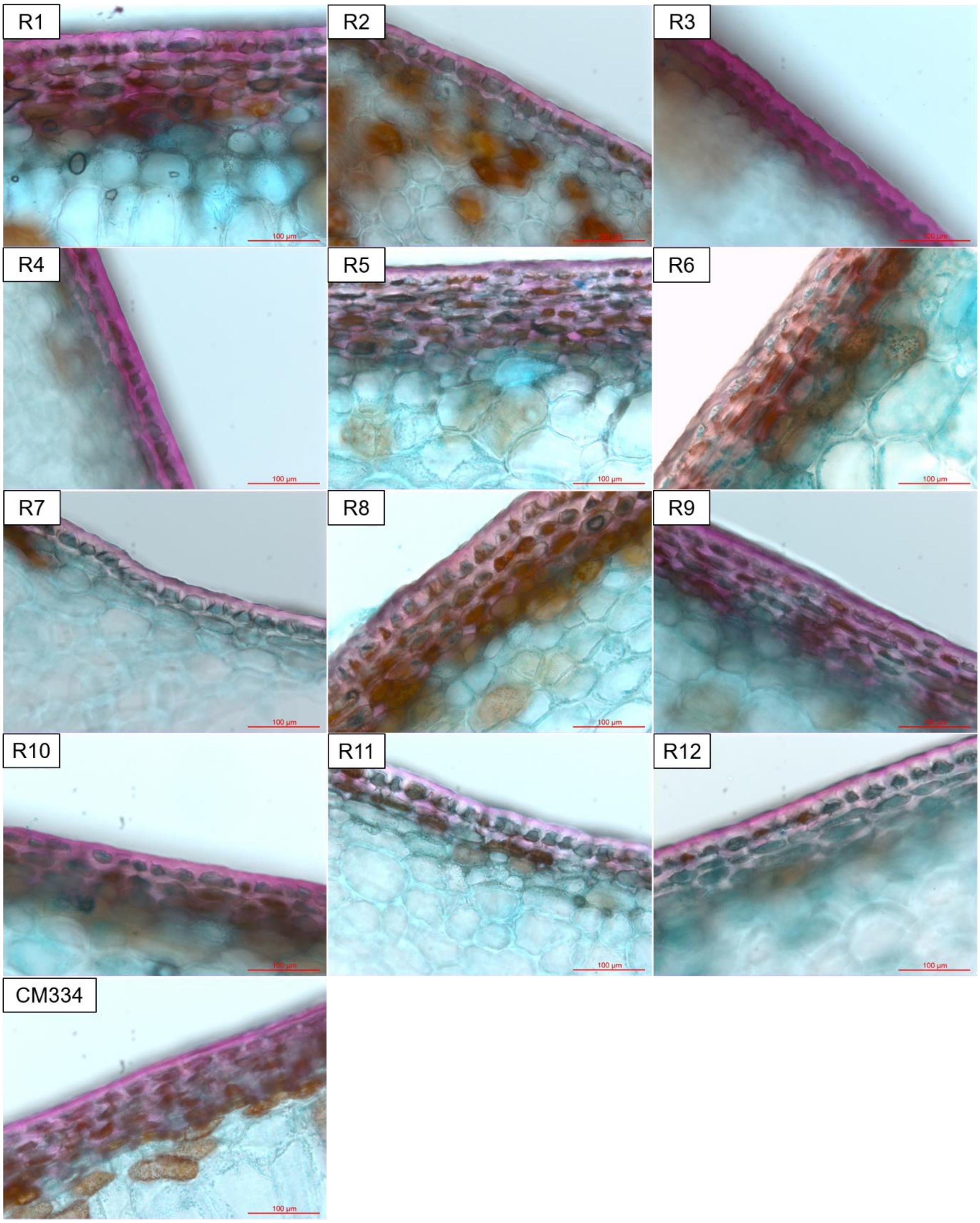

**Fig. S2: Variation in exocarp structure in pepper panel.** Diversity panel, consisting of 13 genotypes, analysed by light microscopy. Exocarp defined as tissue-stained pink using Nile Red stain.

**Table S2: Cutin monomer and cuticular wax components of pepper fruit cuticles.** Cutin monomers and cuticular waxes measured in pepper fruit cuticle discs (1 cm diameter) in mature green and ripe fruits. Values presented represent relative values in comparison with an internal standard. Cutin monomers and cuticular waxes measured in two independent studies. Three biological replicates analysed. Mean ± SEM reported.

| | Mature Green | | | | | Ripe | | | | |
|---|---|---|---|---|---|---|---|---|---|---|
| **Cutin monomer component** | **CM334** | **R3** | **R4** | **R6** | **R8** | **CM334** | **R3** | **R4** | **R6** | **R8** |
| p-Coumaric acid | 5.00 ± 0.10 | 4.47 ± 0.42 | 4.43 ± 0.11 | 5.33 ± 0.34 | 5.06 ± 0.41 | 5.28 ± 0.46 | 3.58 ± 0.64 | 4.47 ± 0.37 | 4.25 ± 0.13 | 4.25 ± 0.95 |
| trans-Ferulic acid | 0.03 ± 0.00 | 0.05 ± 0.01 | 0.11 ± 0.01 | 0.05 ± 0.00 | 0.09 ± 0.02 | 0.04 ± 0.00 | 0.03 ± 0.00 | 0.18 ± 0.02 | 0.15 ± 0.00 | 0.07 ± 0.00 |
| Caffeic acid | 0.04 ± 0.01 | 0.11 ± 0.03 | 0.09 ± 0.01 | 0.04 ± 0.00 | 0.08 ± 0.02 | 0.05 ± 0.01 | 0.03 ± 0.00 | 0.07 ± 0.02 | 0.08 ± 0.01 | 0.06 ± 0.02 |
| Hexadecanoic acid | 0.05 ± 0.00 | 0.08 ± 0.00 | 0.06 ± 0.00 | 0.05 ± 0.00 | 0.05 ± 0.00 | 0.06 ± 0.01 | 0.08 ± 0.01 | 0.07 ± 0.00 | 0.05 ± 0.01 | 0.06 ± 0.02 |
| Hexadecanedioic acid | 0.24 ± 0.01 | 0.25 ± 0.02 | 0.26 ± 0.01 | 0.26 ± 0.01 | 0.54 ± 0.02 | 0.21 ± 0.02 | 0.23 ± 0.04 | 0.18 ± 0.02 | 0.20 ± 0.01 | 0.30 ± 0.04 |
| 6-Hexadecenoic acid | 0.05 ± 0.00 | 0.07 ± 0.01 | 0.05 ± 0.00 | 0.05 ± 0.00 | 0.05 ± 0.00 | 0.06 ± 0.01 | 0.07 ± 0.01 | 0.05 ± 0.00 | 0.05 ± 0.01 | 0.05 ± 0.01 |
| 16-Hydroxyhexadecanoate | 2.00 ± 0.04 | 1.53 ± 0.22 | 4.57 ± 0.29 | 5.66 ± 0.27 | 5.29 ± 0.21 | 1.99 ± 0.13 | 2.03 ± 0.36 | 3.33 ± 0.30 | 3.62 ± 0.19 | 3.38 ± 0.34 |
| 18-hydroxyoctadecanoic acid | 0.03 ± 0.00 | 0.04 ± 0.01 | 0.05 ± 0.00 | 0.03 ± 0.00 | 0.04 ± 0.00 | 0.02 ± 0.01 | 0.04 ± 0.00 | 0.03 ± 0.01 | 0.01 ± 0.00 | 0.02 ± 0.01 |
| Octadeca-9,12-dienoate | 0.01 ± 0.00 | 0.00 ± 0.00 | 0.00 ± 0.00 | 0.00 ± 0.00 | 0.01 ± 0.00 | 0.02 ± 0.00 | 0.01 ± 0.00 | 0.02 ± 0.00 | 0.02 ± 0.00 | 0.02 ± 0.00 |
| 18-hydroxy-9-octadecenoate | 0.21 ± 0.00 | 0.04 ± 0.01 | 0.06 ± 0.01 | 0.08 ± 0.00 | 0.09 ± 0.01 | 0.30 ± 0.02 | 0.17 ± 0.03 | 0.21 ± 0.02 | 0.29 ± 0.02 | 0.22 ± 0.03 |
| 10,16-Dihydroxyhexadecanoic acid | 43.51 ± 6.53 | 17.23 ± 5.95 | 27.94 ± 5.28 | 42.94 ± 7.60 | 58.97 ± 4.85 | 68.97 ± 2.31 | 49.97 ± 7.02 | 51.55 ± 2.95 | 67.70 ± 1.14 | 76.21 ± 6.19 |
| 9,10-epoxy-hydroxyoctadecanoic acid | 1.78 ± 0.38 | 0.59 ± 0.18 | 0.77 ± 0.18 | 1.15 ± 0.28 | 1.44 ± 0.10 | 3.69 ± 0.24 | 1.98 ± 0.25 | 1.76 ± 0.11 | 2.72 ± 0.10 | 3.22 ± 0.73 |
| 9,10-dihydroxyoctadecanedioic acid | 0.28 ± 0.07 | 0.15 ± 0.05 | 0.17 ± 0.05 | 0.25 ± 0.06 | 0.27 ± 0.00 | 0.62 ± 0.03 | 0.57 ± 0.07 | 0.49 ± 0.01 | 0.78 ± 0.00 | 0.95 ± 0.17 |
| 9,10,18-trihydroxyoctadecanoic acid | 0.06 ± 0.01 | 0.03 ± 0.01 | 0.03 ± 0.01 | 0.07 ± 0.02 | 0.09 ± 0.01 | 0.16 ± 0.01 | 0.13 ± 0.02 | 0.13 ± 0.02 | 0.23 ± 0.02 | 0.27 ± 0.06 |
| Total | 53.28 ± 7.16 | 24.65 ± 6.90 | 38.61 ± 5.98 | 55.94 ± 8.59 | 72.05 ± 5.66 | 81.46 ± 3.25 | 58.91 ± 8.46 | 62.55 ± 3.85 | 80.17 ± 1.64 | 89.08 ± 8.56 |

| | Mature Green | | | | | Ripe | | | | |
|---|---|---|---|---|---|---|---|---|---|---|
| **Cuticle wax component** | **CM334** | **R3** | **R4** | **R6** | **R8** | **CM334** | **R3** | **R4** | **R6** | **R8** |
| Tricosane | 0.00 ± 0.00 | 0.01 ± 0.01 | 0.00 ± 0.00 | 0.00 ± 0.00 | 0.01 ± 0.01 | 0.01 ± 0.01 | 0.00 ± 0.00 | 0.00 ± 0.00 | 0.00 ± 0.00 | 0.02 ± 0.02 |
| Tetracosane C2H50 | 0.00 ± 0.00 | 0.04 ± 0.03 | 0.00 ± 0.00 | 0.00 ± 0.00 | 0.01 ± 0.01 | 0.01 ± 0.01 | 0.00 ± 0.00 | 0.00 ± 0.00 | 0.00 ± 0.00 | 0.02 ± 0.02 |
| Pentacosane | 0.01 ± 0.00 | 0.05 ± 0.02 | 0.02 ± 0.00 | 0.01 ± 0.00 | 0.00 ± 0.00 | 0.02 ± 0.01 | 0.02 ± 0.01 | 0.01 ± 0.00 | 0.00 ± 0.00 | 0.01 ± 0.01 |
| Hexacosane | 0.00 ± 0.00 | 0.03 ± 0.02 | 0.01 ± 0.00 | 0.00 ± 0.00 | 0.01 ± 0.00 | 0.01 ± 0.01 | 0.00 ± 0.00 | 0.00 ± 0.00 | 0.00 ± 0.00 | 0.02 ± 0.02 |
| Docosanoic acid | 0.01 ± 0.00 | 0.01 ± 0.00 | 0.02 ± 0.00 | 0.01 ± 0.00 | 0.01 ± 0.00 | 0.01 ± 0.00 | 0.01 ± 0.00 | 0.01 ± 0.00 | 0.01 ± 0.00 | 0.01 ± 0.00 |
| Heptacosane C27H56 | 0.01 ± 0.00 | 0.06 ± 0.03 | 0.04 ± 0.01 | 0.01 ± 0.00 | 0.03 ± 0.01 | 0.02 ± 0.01 | 0.03 ± 0.00 | 0.02 ± 0.00 | 0.01 ± 0.00 | 0.03 ± 0.02 |

| | | | | | | | | | | |
|---|---|---|---|---|---|---|---|---|---|---|
| Octacosane C28H58 | 0.00 ± 0.00 | 0.02 ± 0.02 | 0.00 ± 0.00 | 0.00 ± 0.00 | 0.00 ± 0.00 | 0.00 ± 0.00 | 0.00 ± 0.00 | 0.00 ± 0.00 | 0.00 ± 0.00 | 0.01 ± 0.01 |
| Tetracosanoic acid | 0.02 ± 0.00 | 0.01 ± 0.01 | 0.03 ± 0.01 | 0.03 ± 0.00 | 0.02 ± 0.01 | 0.02 ± 0.01 | 0.02 ± 0.00 | 0.02 ± 0.00 | 0.02 ± 0.00 | 0.02 ± 0.00 |
| Nonacosane C29H60 | 0.01 ± 0.00 | 0.03 ± 0.01 | 0.04 ± 0.01 | 0.03 ± 0.00 | 0.03 ± 0.01 | 0.01 ± 0.01 | 0.01 ± 0.00 | 0.03 ± 0.01 | 0.03 ± 0.00 | 0.03 ± 0.01 |
| Tetratriacontane C34H70 | 0.00 ± 0.00 | 0.00 ± 0.00 | 0.00 ± 0.00 | 0.00 ± 0.00 | 0.00 ± 0.00 | 0.00 ± 0.00 | 0.00 ± 0.00 | 0.00 ± 0.00 | 0.00 ± 0.00 | 0.00 ± 0.00 |
| Hentriacontane | 0.00 ± 0.00 | 0.01 ± 0.00 | 0.02 ± 0.00 | 0.01 ± 0.00 | 0.01 ± 0.00 | 0.00 ± 0.00 | 0.01 ± 0.00 | 0.03 ± 0.01 | 0.01 ± 0.00 | 0.01 ± 0.00 |
| β-Sitosterol | 0.01 ± 0.00 | 0.00 ± 0.00 | 0.01 ± 0.00 | 0.01 ± 0.00 | 0.00 ± 0.00 | 0.01 ± 0.00 | 0.00 ± 0.00 | 0.01 ± 0.00 | 0.01 ± 0.00 | 0.01 ± 0.00 |
| β-amyrin | 0.05 ± 0.01 | 0.03 ± 0.01 | 0.03 ± 0.01 | 0.02 ± 0.00 | 0.03 ± 0.00 | 0.07 ± 0.03 | 0.02 ± 0.00 | 0.04 ± 0.01 | 0.02 ± 0.00 | 0.03 ± 0.01 |
| α-amyrin | 0.05 ± 0.01 | 0.03 ± 0.01 | 0.03 ± 0.01 | 0.02 ± 0.00 | 0.03 ± 0.00 | 0.07 ± 0.03 | 0.02 ± 0.00 | 0.04 ± 0.01 | 0.02 ± 0.00 | 0.03 ± 0.01 |
| Lupeol | 0.03 ± 0.01 | 0.02 ± 0.01 | 0.02 ± 0.00 | 0.01 ± 0.00 | 0.02 ± 0.00 | 0.04 ± 0.02 | 0.01 ± 0.00 | 0.02 ± 0.00 | 0.01 ± 0.00 | 0.02 ± 0.00 |
| Total | 0.21 ± 0.04 | 0.36 ± 0.13 | 0.27 ± 0.03 | 0.16 ± 0.01 | 0.20 ± 0.04 | 0.31 ± 0.17 | 0.16 ± 0.02 | 0.24 ± 0.05 | 0.14 ± 0.00 | 0.26 ± 0.14 |

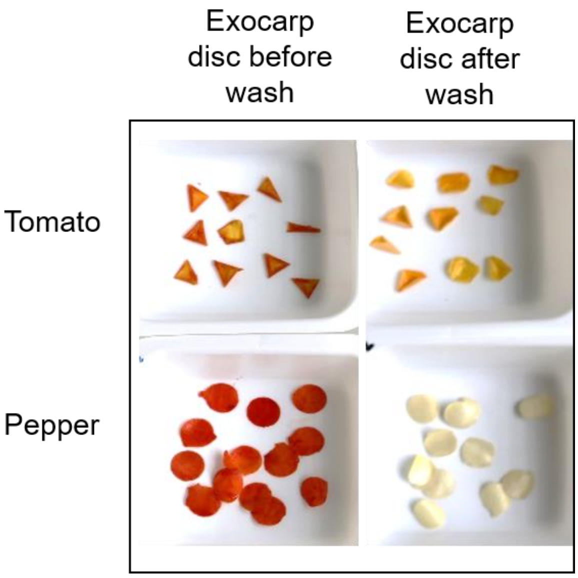

**Fig. S3: Isolated exocarp discs of pepper and tomato fruits, before and after chloroform washing.**
